# Tumor-intrinsic LKB1-LIF signaling axis establishes a myeloid niche to promote immune evasion and tumor growth

**DOI:** 10.1101/2023.07.15.549147

**Authors:** Ali Rashidfarrokhi, Ray Pillai, Yuan Hao, Warren L. Wu, Burcu Karadal-Ferrena, Sofia G. Dimitriadoy, Michael Cross, Anna H. Yeaton, Shih Ming Huang, Arjun J. Bhutkar, Alberto Herrera, Sahith Rajalingam, Makiko Hayashi, Kuan-lin Huang, Eric Bartnicki, Anastasia-Maria Zavitsanou, Corrin A. Wohlhieter, Sarah E. Leboeuf, Ting Chen, Cynthia Loomis, Valeria Mezzano, Ruth Kulicke, Fred P. Davis, Nicolas Stransky, Gromoslaw A. Smolen, Charles M. Rudin, Andre L. Moreira, Kamal M. Khanna, Harvey I. Pass, Kwok-Kin Wong, Shohei Koide, Aristotelis Tsirigos, Sergei B. Koralov, Thales Papagiannakopoulos

## Abstract

Tumor mutations can influence the surrounding microenvironment leading to suppression of anti-tumor immune responses and thereby contributing to tumor progression and failure of cancer therapies. Here we use genetically engineered lung cancer mouse models and patient samples to dissect how *LKB1* mutations accelerate tumor growth by reshaping the immune microenvironment. Comprehensive immune profiling of *LKB1*-mutant vs wildtype tumors revealed dramatic changes in myeloid cells, specifically enrichment of Arg1^+^ interstitial macrophages and SiglecF^Hi^ neutrophils. We discovered a novel mechanism whereby autocrine LIF signaling in *Lkb1*-mutant tumors drives tumorigenesis by reprogramming myeloid cells in the immune microenvironment. Inhibiting LIF signaling in *Lkb1*-mutant tumors, via gene targeting or with a neutralizing antibody, resulted in a striking reduction in Arg1^+^ interstitial macrophages and SiglecF^Hi^ neutrophils, expansion of antigen specific T cells, and inhibition of tumor progression. Thus, targeting LIF signaling provides a new therapeutic approach to reverse the immunosuppressive microenvironment of *LKB1*-mutant tumors.

## INTRODUCTION

Greater understanding of tumor cell interactions with the tumor immune microenvironment (TIME) is driving the rapid evolution of therapeutic strategies for cancer. Lung cancer is the leading cause of cancer related deaths worldwide with lung adenocarcinoma (LUAD) being the most common histologic type of lung cancer (Lareau et al., 2021). Currently the primary treatment modality for advanced LUAD utilizes immune checkpoint inhibitors (ICI) to augment anti-tumor immune responses and inhibit tumor progression (Gandhi et al., 2018). Despite the widespread use of ICI in LUAD, the overall response rates remain low (Jeanson et al., 2019). The genetic heterogeneity of tumors likely contributes to the poor responses to ICI. While some driver gene mutations are known to sensitize tumors to specific targeted therapies, many mutations induce resistance to ICI through unknown mechanisms. Mutational inactivation of Liver kinase B1 (*LKB1;* also known as serine/threonine kinase 11 *STK11*) is among the most frequent genetic aberrations occurring in about 20% of patients with LUAD and this mutation often co-occurs with loss-of-function mutations in Kelch like ECH associated protein 1 (*KEAP1)* (Arbour et al., 2018; Best et al., 2019; Cancer Genome Atlas Research, 2014; Papillon-Cavanagh et al., 2020; Wohlhieter et al., 2020). These mutations frequently co-occur with *KRAS* mutations and are associated with poor patient outcomes due to resistance to current standard of care treatments including chemotherapy combined with ICI as well as the newly developed KRAS G12C inhibitors (Arbour *et al*., 2018; Cristescu et al., 2018; Papillon-Cavanagh *et al*., 2020; Ricciuti et al., 2020; Shen et al., 2019; Skoulidis et al., 2021; Wohlhieter *et al*., 2020). Due to the lack of effective treatment options, understanding how *LKB1*-mutant and *KEAP1*-mutant tumors alter the TIME and affect ICI efficacy in LUAD is essential.

Inflammation either in the form of chronic inflammatory disease or as the byproduct of tumor-derived inflammation can greatly impact the function of immune cells (Greten and Grivennikov, 2019). Specifically, inflammation can change the plasticity and heterogeneity of both tumor cells and the surrounding TIME (Grivennikov et al., 2010). The field of tumor immunology research has largely focused on adaptive immune responses and the tumor-intrinsic mechanisms of evading lymphocytes (Nguyen and Spranger, 2020; Spranger and Gajewski, 2018). However, the predominant immune cells in the lung are myeloid populations which play a significant role in lung inflammation in the context of both infection and cancer (Binnewies et al., 2018; Wellenstein and de Visser, 2018). Myeloid cells play an integral role in both activating T cells and regulating tumor growth. However, the composition of myeloid cell populations, within tumors, specifically macrophages and neutrophils, and how those cells are impacted by intrinsic tumor mutations has largely been unexplored in LUAD (Casanova-Acebes et al., 2021; Engblom et al., 2017; Pfirschke et al., 2020). *In vitro* models have been developed to understand the impact of macrophages and neutrophils on T cell function, but these models fail to capture the complexities of the TIME and do not address how myeloid cells modulate anti-tumor T cell responses *in vivo*.

In this study we examine the role of tumor specific *Lkb1-*mutations in altering lung inflammation using genetically engineered mouse models (GEMMs) of LUAD (Best *et al*., 2019; Hollstein et al., 2019; Koyama et al., 2016a; Murray et al., 2019; Romero et al., 2017; Sánchez-Rivera et al., 2014). By utilizing multiple complementary modalities including flow cytometry, multi-color immunofluorescence (multi-IF), and single cell RNA sequencing (scRNA-seq) we found that tumor-intrinsic *Lkb1*-mutations reshape the TIME, driving a reduction in levels of alveolar macrophages with concomitant increase in SiglecF^Hi^ neutrophils and Arg1^+^ interstitial macrophages along with augmented expression of pro-inflammatory cytokines and chemokines. Mechanistically, we discovered that *Lkb1*-mutant tumors upregulate expression of the cytokine Leukemia inhibitory factor (LIF), which signals in an autocrine manner through its receptor, LIFR, on tumor cells to activate an inflammatory signaling cascade driving the altered myeloid composition and transcriptional program of the TIME. Genetic knockout or antibody-based neutralization of LIF signaling reversed the myeloid cell infiltration and inflammatory phenotype. As a result of LIF blockade, myeloid changes in the TIME were accompanied by enhanced T cell clonal expansion and restrained tumor growth. These findings suggest that *LKB1*-mutant lung tumors generate a LIF-dependent inflammatory response associated with immunosuppressive myeloid cells that can inhibit T cell function and promote tumor growth.

## RESULTS

### Tumor intrinsic *Lkb1* mutations alter myeloid landscape in the TIME

To investigate the role of *Lkb1-* and *Keap1*-mutations in altering the TIME and determine how these tumor mutations impact anti-tumor immune responses we established GEMMs that recapitulate human LUAD. We utilized a two paired guide RNA CRISPR/Cas9 somatic editing system combined with the *Kras*^LSL-G12D/+^ p53^fl/fl^ (KP) GEMMs (Vidigal and Ventura, 2015) that enables us to investigate the impact of *Lkb1*- and *Keap1*-mutations individually and in combination in the context of *Kras;p53-*mutant (*Kras*^G12D/+^*; p53*^-/-^) lung cancer (Figure 1A, Figure S1A) (Ding et al., 2021; Platt et al., 2014; Romero *et al*., 2017; Sánchez-Rivera *et al*., 2014). We monitored tumor growth by MRI and collected tumor bearing lungs for histological analyses and immune profiling at 6- and 11-weeks post tumor initiation (Figure 1A). Consistent with prior work, we observed that *Lkb1*-mutant tumors are significantly more aggressive compared to *Lkb1*-wildtype tumors based on increased tumor burden as seen by MRI 11 weeks after infection (Figure 1B) (Hollstein *et al*., 2019; Koyama *et al*., 2016a; Murray *et al*., 2019). Interestingly, concurrent mutation of *Keap1* and *Lkb1* did not result in augmented tumor burden compared to tumors with only *Lkb1* mutation, despite both mutations individually contributing to tumor progression (Figure S1B and S1C). However, *Lkb1*-*Keap1* co-mutated tumors did have the highest tumor grade (Figure S1D).

**Figure 1:**
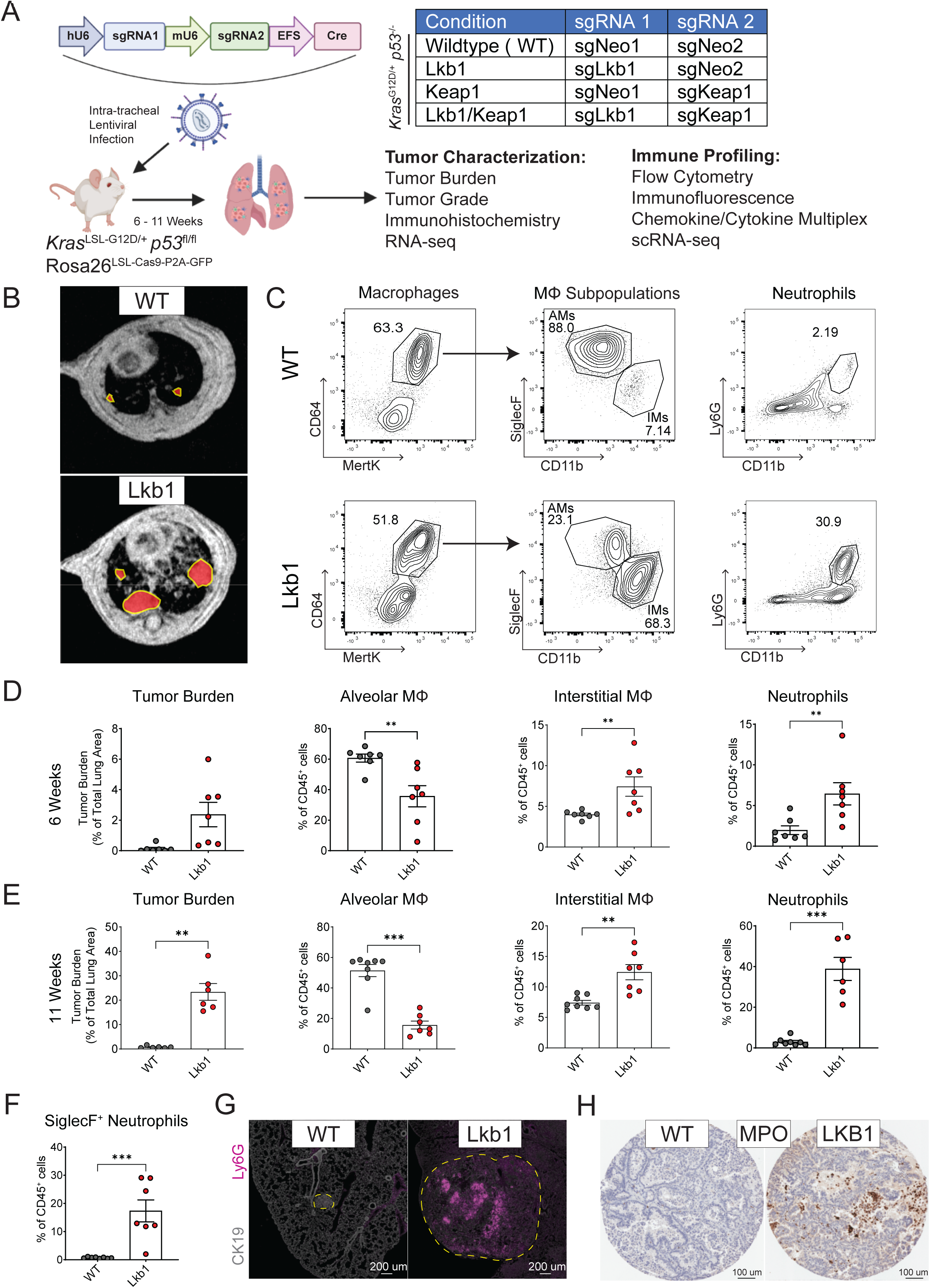
*Lkb1*-mutations alter myeloid cell composition in the TIME. (A) Schematic representation of autochthonous lung tumors generated using intra-tracheal lentivirus delivery with dual guide CRISPR/Cas9 editing in *Kras*^LSL-G12D/+^ *p53*^fl/fl^ Rosa26^LSL-Cas9-P2A-GFP^ mice. Four genetic conditions are generated with guide RNAs indicated in the table. (B) Representative MRI images of mouse lungs with wildtype or *Lkb1*-mutant tumors 11 weeks post infection. Tumors are outlined in yellow and highlighted in red. (C) Representative flow cytometry plots of neutrophil and macrophage subpopulations in wildtype (WT) and *Lkb1*-mutant (Lkb1) tumor bearing lungs. Alveolar macrophages (AMs) and interstitial macrophages (IMs) are identified as distinct macrophage (Mϕ) subsets. (D, E) Bar plot of tumor burden as a percent of cross-sectional tumor area from total lung area measured on a midline H&E section at (D) 6 weeks and (E) 11 weeks post tumor initiation. Myeloid cell infiltration (alveolar macrophages, interstitial macrophages, and neutrophils) represented as a percentage of total tissue infiltrating immune cells (CD45^+^ non-circulating/circulating CD45^-^) in the indicated genotype for each timepoint. (F) SiglecF^+^ Neutrophil infiltration represented as a percentage of total tissue infiltrating immune cells (CD45^+^ CD45-Cir^-^) in each indicated genotype. (G) Immunofluorescence images of tumor bearing lungs showing Ly6G (purple) and CK19 (grey) staining. Tumor is outlined in yellow. Scale bar, 200 μm. (H) Representative image of myeloperoxidase (MPO) staining in a human tumor microarray. Scale bar, 100 μm. Results are displayed as mean ± SEM. n of 6 - 8 mice were used for each mouse experiment. Statistical analysis was performed using Mann Whitney U or one-way ANOVA with Tukey’s test where appropriate. * p< 0.05 ** p<0.01 *** p<0.001 **** p< 0.0001.

Innate immune cells in the TIME, including macrophages and neutrophils, can influence cancer progression and immune evasion (Engblom et al., 2016; Lavin et al., 2015). Despite the important role of innate cells in shaping immune responses, the diversity and function of macrophages and neutrophils in genetic subsets of LUAD has not been well-defined. To characterize the immune landscape of *Lkb1*-mutant LUAD GEMMs we used multicolor flow cytometry (representative flow cytometry plots and gating shown in Figure 1C and S1E) at 6 weeks post tumor initiation to compare immune infiltration during early stages of tumor growth (Figure 1D) and 11 weeks post initiation where *Lkb1*-mutant tumors display higher tumor burden (Figure 1E). Flow cytometry analysis of tumor bearing lungs revealed a significant decrease in alveolar macrophages (AMs) and an increase in interstitial macrophages (IMs) in *Lkb1*-mutant tumors (Figure 1C, 1D, 1E, S1F). Consistent with prior studies, *Lkb1*-mutant tumors have a dramatic increase in neutrophils (Figure 1C, 1D, 1E, S1F) (Koyama *et al*., 2016a). Notably, these same changes in myeloid populations are seen at 6 weeks (Figure 1D) after tumor initiation where there is no significant difference in tumor burden compared to *Lkb1*-WT tumors suggestive that these changes are related to tumor genetics rather than tumor burden. Further supporting this, at 11 weeks post infection mice bearing *Keap1*-mutant tumors have significantly more tumor burden than WT controls yet do not show any significant changes in myeloid populations (Figure S1B, S1G). In addition, our profiling revealed that the innate immune infiltrate of *Lkb1*-mutant and *Lkb1*/*Keap1* co-mutant tumors have a very similar immune phenotype (Figure S1G), suggesting that it is specifically mutations in *Lkb1* that are responsible for the observed changes in the TIME. Therefore, we focused on the immune microenvironment of *Lkb1*-mutant tumors for our subsequent analysis.

Neutrophils are thought to have both pro- and anti-tumorigenic roles (Coffelt et al., 2016). Recently, pro-tumorigenic SiglecF^+^ neutrophils have been identified to infiltrate lungs in transplant models of lung adenocarcinoma (Engblom *et al*., 2017; Pfirschke *et al*., 2020). Because these SiglecF^+^ neutrophils promote tumorigenesis, we specifically looked for them in the *Lkb1*-mutant GEMM. Strikingly, SiglecF^+^ neutrophils were significantly enriched in the setting of *Lkb1*-mutant tumors increasing from ∼ 1% to ∼ 18% of immune cells (Figure 1F). Immunofluorescent staining for Ly6G confirmed that neutrophils are increased in *Lkb1*-mutant tumors (Figure 1G) consistent with the aforementioned data. Next, we sought to validate these mouse model findings in LUAD patient samples. Using a genetically defined tumor microarray we looked at tumor cores from patients and stained for myeloperoxidase (MPO) as a marker for neutrophils by immunohistochemistry (IHC). Similar to the mouse immunofluorescence, *LKB1*-mutant patient tumor cores had increased neutrophils further supporting that loss of function mutations in *LKB1* tumors are a driver for changes in neutrophil infiltration (Figure 1H, S1H).

We hypothesized that *Lkb1*-mutant tumor inflammatory pattern driven by changes in macrophage and neutrophil populations are creating an immunosuppressive microenvironment that promotes escape from antitumor immune responses. Since T cells play a critical role in anti-tumor immune responses, we examined effector function of T cells isolated from *Lkb1*-mutant tumors. Stimulation of *ex vivo* T cells collected from the lungs of animals with *Lkb1*-mutant tumors revealed dramatic reduction in IFNγ and TNFα production compared to T cells isolated from control tumors, indicating that T cell effector function is severely impaired in *Lkb1*-mutant tumors (Figure S1I and S1J). collectively, these findings suggest that *Lkb1*-mutant tumors induce an inflammatory lung microenvironment characterized by increased IMs and neutrophils as well as reduced T cell effector function.

### *Lkb1-*mutant tumors reprogram myeloid cells in the TIME

Given the dramatic changes within the myeloid subsets of *Lkb1-*mutant tumors, we hypothesized that these immune cells are promoting an immunosuppressive TIME. To obtain a more granular view of the immunosuppressive TIME in *Lkb1*-mutant lung tumors, we performed single cell RNA-seq profiling of live extravascular CD45^+^ cells from whole lung digests. Unsupervised clustering analysis enabled identification of different immune populations which were annotated based on gene expression signatures (Figure S2A, S2B and S2C). To fully elucidate the impact of tumor intrinsic *Lkb1* loss on macrophage and neutrophil populations within TIME, we sub-clustered macrophage and neutrophil populations into 9 and 8 unique clusters, respectively. Macrophage (Mϕ) sub-clustering revealed 4 clusters of AMs: alveolar Mϕ1, cycling Mϕ, ISG^Hi^ Mϕ, Spp1^+^ Mϕ1 and 5 clusters of IMs: Interstitial Mϕ1, Monocyte derived Mϕ (mo-Mϕ), CX3CR1^+^ Mϕ, Lyve1^+^ Mϕ, and Spp1^+^ Mϕ2 (Figure 2A, 2B, and S2D). In agreement with our flow cytometry data, there is a dramatic increase in the proportion of IMs with a reduction of AMs in the setting of *Lkb1*-mutant tumors (Figure 2B).

**Figure 2:**
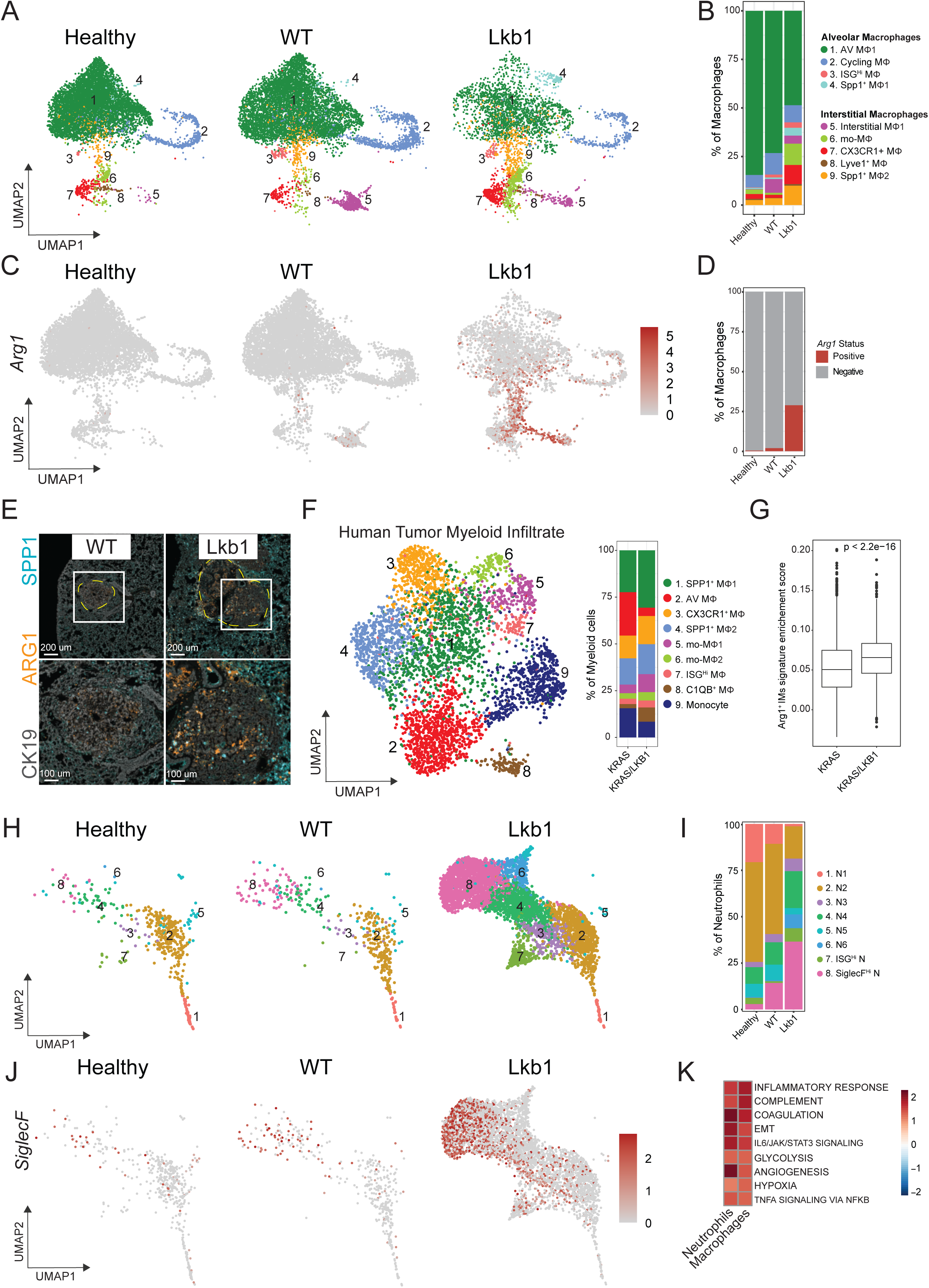
*Lkb1*-mutant tumors influence myeloid cell transcriptional program, inducing an immunosuppressive phenotype. (A) UMAP plots of macrophage sub-clusters in Healthy lungs, *Lkb1*-WT, and *Lkb1*-mutant tumors. Identity of each cluster is based on gene expression. (B) Quantification of macrophage sub-clusters seen in (A) by different conditions. Clusters are separated into alveolar and interstitial macrophages. (C) UMAP plots of macrophage syv-clusters showing *Arg1* expression in Healthy lungs, *Lkb1*-WT, and *Lkb1*-mutant tumors. (D) Quantification of *Arg1+* macrophages seen in (C) expressed as a percentage of total macrophages. (E) Multi-immunofluorescence images showing ARG1 (orange), SPP1 (cyan), and CK19 (grey) staining in WT and *Lkb1*-mutant tumors. Scale bar, 200 μm (left images) and 100 μm (right images). (F) UMAP of myeloid cells analyzed by single-nuclei RNA-seq of human LUAD samples. Quantification of myeloid cell subclusters is shown stratified by mutation status: *KRAS* (n=14), *KRAS LKB1* (n=4). (G) Enrichment score for Arg1^+^ interstitial macrophages stratified by *LKB1*-mutation status. Wilcoxon rank sum test was used to analyze the data. (H) UMAP plots of neutrophil sub-clusters comparing subpopulations in lungs of healthy mice compared to mice with wildtype and *Lkb1*-mutant tumors. (I) Quantification of neutrophil sub-clusters seen in panel (H) by different conditions. (J) UMAP plots of *SiglecF* expression in neutrophils shown in panel (H) (K) Significantly enriched pathways (FDR <0.1) in macrophages and neutrophils comparing myeloid cells from *Lkb1*-mutant tumors to WT lung tumors.

We next evaluated for the immunosuppressive potential of these myeloid cells isolated from tumors of different genotypes. One well characterized immunosuppressive marker, Arginase 1 (*Arg1*), was dramatically upregulated in macrophages isolated from *Lkb1*-mutant tumors (Figure 2C, 2D, and S2E). ARG1 is one of the hallmarks of immunosuppressive myeloid cells and involved in suppressing T cell function and proliferation (Bronte et al., 2003; Fu et al., 2022; Geiger et al., 2016; Katzenelenbogen et al., 2020; Miret et al., 2019; Rodriguez et al., 2004). In our single cell RNA-seq dataset, *Arg1* was predominantly increased in IMs rather than AMs suggestive that these IMs are the key macrophage population driving the immunosuppressive microenvironment. We also observed the emergence of two Spp1+ clusters marked by increased *Spp1* expression in *Lkb1*-mutant tumors (Figure 2B and S2F). Although the function of *Spp1* is generally unknown in cancer models, infiltration of SPP1^+^ macrophages are associated with tumor progression and resistance to immunotherapy (Cheng et al., 2021; Qi et al., 2022; Zhang et al., 2020). To validate the transcriptional findings from the single cell RNA-seq data, we performed multi-IF staining for ARG1 and SPP1 which confirmed increased protein expression of these markers in sections from lungs bearing *Lkb1*-mutant tumors (Figure 2E).

In order to determine whether similar changes in macrophage populations are also observed in patients, we performed single-nucleus RNA-seq on immune cells from *LKB1-* mutant and *LKB1*-WT human LUAD tumors (Figure S3A). Consistent with the GEMM data, we found an increase in Spp1*^+^* macrophage cluster and a reduction in AMs in *LKB1-* mutant tumors (Figure 2F and S3B). Furthermore, we sought to determine whether the immunosuppressive IMs seen in *Lkb1*-mutant GEMMs are also present in *LKB1*-mutant LUAD patient tumors. Using gene expression data from Arg1^+^ IMs in *Lkb1*-mutant lung mouse tumors we created an Arg1^+^ IM gene signature (Figure S3C). We found that this Arg1^+^ IM signature was enriched in macrophage populations from human *LKB1*-mutant LUAD (Figure 2G). Furthermore, using a bulk RNA-seq dataset from the TCGA we found that this Arg1^+^ signature is associated with worst survival (Figure S3D). Taken together, these data suggest that IMs expressing immunosuppressive signatures are enriched in *LKB1*-mutant tumors and may play an important role in promoting immune evasion and tumor growth.

Finally, we sub-clustered the neutrophil population to 8 subclusters: N1, N2, N3. N4, N5, N6, ISG^Hi^ N, and SiglecF^Hi^ N (Figure 2H and S4A). Overall, we observed that neutrophils are enriched in the *Lkb1*-mutant condition (Figure 2H). In addition, *Lkb1*-mutant tumor bearing mice had an increased proportion of clusters N4, N6, ISG^Hi^ N, and SiglecF^Hi^ N (Figure 2I, 2J, and S4B), the latter of which was also demonstrated by flow cytometry (Figure 1F). The neutrophils from *Lkb1*-mutant tumors were found to upregulate a series of genes including *Tnf, Vegfa, Hif1a, Havcr2, Fcgr2b, Csf1* and *Ccl3* (Figure S4C), transcripts that have been previously implicated in tumor promoting neutrophil populations that drive tumor proliferation, angiogenesis, T cell suppression, and macrophage recruitment (Engblom *et al*., 2017).

To determine the pathways that are altered in macrophages and neutrophils, we compared differentially expressed genes (DEGs) in *Lkb1-*mutant to WT tumors. We observed an enrichment of similar pathways in both neutrophils and macrophages involved in suppression of anti-tumor immunity including IL6/JAK/STAT3 signaling, hypoxia, inflammatory response, and TNFA signaling (Arts et al., 2016; Doedens et al., 2010; Engblom *et al*., 2016; Kortylewski et al., 2005; Movahedi et al., 2010; Vitale et al., 2019; Yu et al., 2007) (Figure 2K). Collectively, our single cell analysis of mouse and human tumors confirms that *LKB1*-mutations dramatically impact the infiltration and transcriptional program of macrophages and neutrophils, creating an immunosuppressive microenvironment.

### *Lkb1*-mutant tumors produce leukemia inhibitory factor

Since similar pathways were upregulated in tumor-infiltrating neutrophils and macrophages within *Lkb1*-mutant tumors, we suspected that a common factor is inducing these transcriptional programs. To determine how *Lkb1*-mutant tumors were altering the infiltration and transcriptional state of myeloid cells in the TIME, we sorted *Lkb1*-mutant and WT tumor cells from the GEMMs and performed bulk RNA-seq (Figure S5A). Pathway enrichment analysis demonstrated upregulated of pathways in the malignant cells similar to those seen in macrophages and neutrophils, including inflammatory response, TNFA signaling, and IL6/JAK/STAT3 signaling (Figure 3A and S5B). Based on the observation that both tumor cells and myeloid cells express many of the same pathways related to JAK-STAT3 signaling and inflammatory pathways, we hypothesized that common pro-inflammatory mediators were driving the reprogramming of the TIME. Therefore, we decided to look at chemokines and cytokines specifically upregulated in *Lkb1*-mutant tumors. Analysis of our RNA-seq dataset revealed that *Lkb1*-mutant tumors had increased expression of known neutrophil chemoattractants such as *Cxcl1*, *Cxcl3*, *Cxcl5* as well as cytokines such as *Il1a*, *Il6*, *Csf3*, and *Il33* in agreement with previous studies (Hollstein *et al*., 2019; Koyama *et al*., 2016a; Murray *et al*., 2019; Quail et al., 2022) (Figure 3B). In addition to this, we discovered that the cytokine leukemia inhibitory factor (*Lif)* was upregulated in *Lkb1*-mutant tumors (Figure 3B). LIF plays an important role in stem cell self-renewal and regulation of differentiation (He et al., 2006; Mathieu et al., 2012; Rathjen et al., 1990; Stewart et al., 1992a) and has recently been implicated in promoting cancer cell growth (Albrengues et al., 2014; Liu et al., 2013; Pascual-Garcia et al., 2019; Penuelas et al., 2009; Shi et al., 2019). However, its role in lung cancer and specifically in the regulation of tumor-elicited inflammation had not been previously described. Therefore, we focused on characterizing whether LIF signaling plays a functional role in reshaping the TIME in *Lkb1*-mutant tumors.

**Figure 3:**
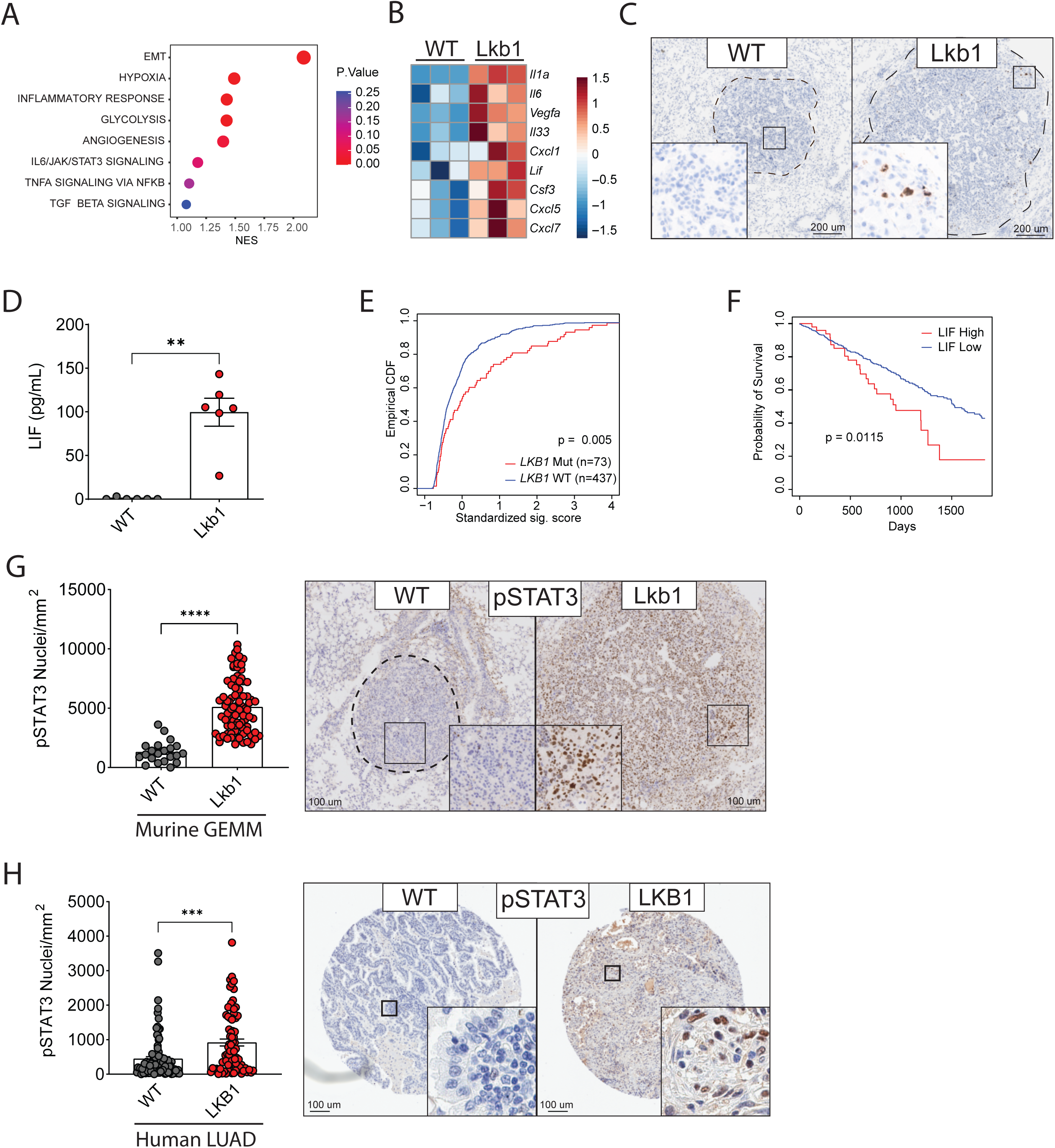
Tumor-intrinsic *Lkb1* loss induces *Lif* expression that contributes to increased STAT3 signaling. (A) Significantly enriched transcriptional pathways in *Lkb1*-mutant tumor cells compared to wildtype tumors sorted from lung cancer mouse model revealed by RNA-seq analysis. (B) Gene expression heatmap of selected significant genes (*Il1a, Il6*, Vegfa, *Il33*, *Cxcl1, Lif, Csf3, Cxcl6, Cxcl7*) (p< 0.05) measured by RNA-seq on sorted WT and *Lkb1-*mutant tumors. (C) *In situ* RNA hybridization using RNA scope showing *Lif* expression in lung sections from WT and *Lkb1*-mutant tumors. Tumors are outlined in black. Scale bar, 200 μm. (D) LIF protein levels in bronchoalveolar lavage fluid of tumor bearing mice of each genotype measured by chemokine/cytokine multiplex assay. (E) Empirical cumulative distribution function (CDF) plots showing *LIF* expression for TCGA LUAD patients grouped by *LKB1* mutation status. (F) Kaplan–Meier 5-year survival curves comparing patients in the TCGA LUAD cohort (G) Quantification of positive intra-tumoral pSTAT3 staining by immunohistochemistry in a human lung tumor microarray. Representative images of a wildtype (WT) and *LKB1* mutant (LKB1) tumor are shown. Scale bar, 100 μm. (H) Kaplan–Meier 5-year survival curves comparing patients in the TCGA LUAD cohort stratified by either high (top 10%) or low (rest of the cohort) *LIF* expression. Results for LIF protein levels are displayed as mean ± SEM. n of 6 mice were used for each mouse experiment. For pSTAT3 analysis individual mouse tumors or human tumor cores are displayed. Statistical analysis was performed using Mann Whitney U or one-way ANOVA with Tukey’s test where appropriate. * p< 0.05 ** p<0.01 *** p<0.001 **** p< 0.0001.

To confirm the augmented expression of *Lif* in *Lkb1*-mutant tumors, we utilized two approaches: 1) RNA *in situ* hybridization to verify that *Lif* is expressed intratumorally in the GEMMs (Figure 3C) and 2) multiplex chemokine/cytokine array to evaluate LIF levels in the bronchoalveolar lavage (BAL) fluid of tumor-bearing GEMMs (Figure 3D and S5C). In order to demonstrate that *Lkb1* loss alone was sufficient to induce expression of *Lif*, we generated murine lung and pancreatic cancer KP cells deficient for *Lkb1*. As expected, we observed increased expression of *Lif* in both *Lkb1* knockout (KO) cells confirming that LKB1 regulates the transcription of *Lif* in multiple cancer types (Figure S5D). To validate that *Lif* is similarly upregulated in *LKB1-*mutant human tumors, we analyzed bulk RNA-seq TCGA dataset. Our analysis showed that *LKB1*-mutant LUAD have increased *LIF* expression compared to *LKB1* wildtype tumors (Figure 3E). Furthermore, survival analysis of TCGA LUAD data revealed that patients with high *LIF* expression have significantly worse survival (Figure 3F).

Since JAK-STAT signaling was upregulated in *Lkb1*-mutants (Figure 3A), and STAT3 phosphorylation is known to be downstream of LIFR (Nicola and Babon, 2015; Stahl et al., 1994; Taga and Kishimoto, 1997), we hypothesized that LIF was signaling in an autocrine manner. Elevated pSTAT3, the transcriptionally active form of STAT3, was observed in *Lkb1*-mutant tumors upon IHC staining supporting the idea of autocrine LIF signaling in this mouse model (Figure 3G). Furthermore, analysis of the TMA also demonstrated increased pSTAT3 specifically in *LKB1*-mutant tumor cores (Figure 3H) consistent with our lung cancer GEMM. Together, our results suggest that *LKB1* loss leads to increased LIF production, and that this contributes to pro-inflammatory JAK-STAT signaling within the tumor cells via autocrine engagement of the LIFR, the receptor for LIF.

### Autocrine LIF signaling establishes an inflammatory niche that supports tumor growth

Given that *Lkb1*-mutant tumors demonstrated increased LIF production and pSTAT3 levels, we hypothesized that LIF was acting through an autocrine mechanism to upregulate cytokines and chemokines generating a pro-inflammatory TIME (Figure 4A). To investigate this hypothesis, we used somatic CRISPR/Cas9 editing to knockout *Lif* or its receptor *Lifr* in the setting of *Lkb1*-mutation in KP GEMMs to assess the impact of LIF signaling on tumor progression and the TIME (Figure 4B and S6A). Both *Lif* and *Lifr* KO significantly impaired tumor growth as evidenced by reduced tumor burden quantified by MRI at 11 weeks (Figure 4C and 4D), suggesting that LIF signaling is active in *Lkb1*-mutant tumors and supports tumor progression. Consistent with this reduction in tumor burden, KO of *Lifr* also improved survival of mice with *Kras*-driven *p53*^+/+^ *Lkb1*-mutant, demonstrating that the significance of LIF signaling is not restricted to a KP model (Figure S6B). To evaluate the tumor intrinsic effects of LIF signaling on tumor growth, we measured *in vitro* growth of *Lif* KO and *Lifr* KO cell lines and observed no impact on proliferation (Figure S6C), suggestive that the differences in tumor burden and survival found in the GEMMs is dependent on tumor extrinsic factors such as immune surveillance. We validated KO of *Lif* in our *Lkb1*-mutant lung cancer GEMMs and found nearly undetectable levels of LIF in the *Lif* KO condition (Figure 4E), demonstrating that the majority of LIF found in the BAL is produced by *Lkb1*-mutant malignant cells and rather than other cells within the tumor microenvironment. Interestingly, KO of *Lifr* had comparable levels of LIF to controls despite having significantly less burden (Figure 4E). Overall, these findings suggest that LIF promotes tumor growth by autocrine signaling through the LIFR on tumor cells. Furthermore, levels of other cytokines found to be elevated within *Lkb1*-mutant TIME, such as IL-6 and CSF3, were significantly reduced in both the *Lif* and *Lifr* KO conditions (Figure S6D and S6E). Since LIF primarily signals through pSTAT3, we confirmed that pSTAT3 was reduced in both *Lif* KO and *Lifr* KO tumors (Figure 4F and S6F).

**Figure 4:**
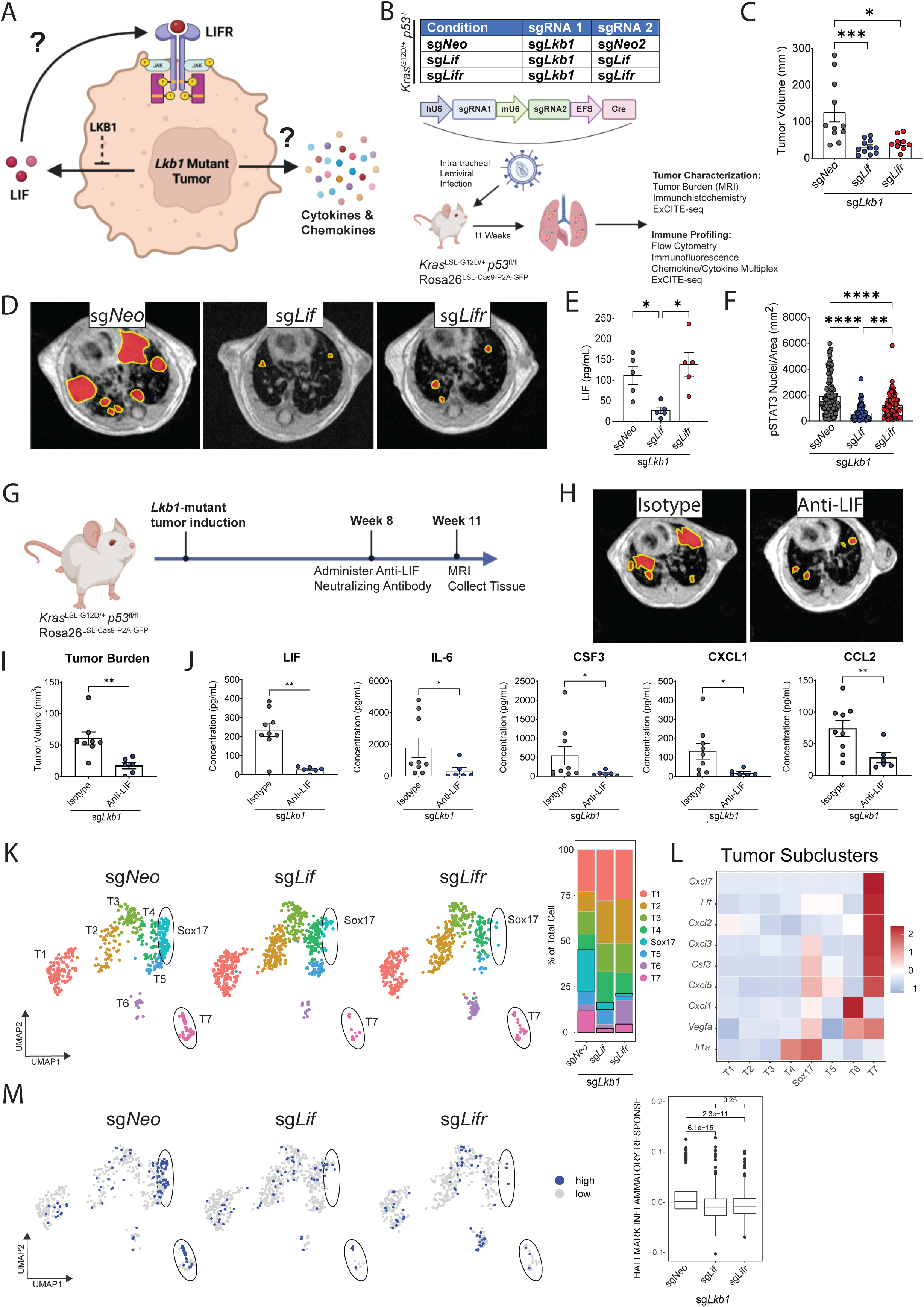
Tumor intrinsic LIF/LIFR signaling generates an inflammatory niche and promotes tumor growth. (A) Working model of *Lkb1*-mutant tumors induction of autocrine LIF signaling and inflammatory cytokines and chemokines. (B) Schematic representation of our autochthonous lung tumors generated using intra-tracheal lentivirus delivery with dual guide CRISPR/Cas9 editing in *Kras*^LSL-G12D/+^ *p53*^fl/fl^ Rosa26^LSL-Cas9-P2A-GFP^ mice to generate *Lkb1*-mutant tumors with knockout of *Lif* (sg*Lif*) or *Lifr (*sg*Lifr)*. (C) *Lkb1*-mutant lung tumors were initiated with knockout of *Lif*, *Lifr*, or control (sgNeo). Tumor burden quantified by MRI. (D) Representative images for each genotype from (C) is shown. Tumors are highlighted in red and outlined in yellow. (E) LIF concentration in bronchoalveolar lavage fluid of *Lkb1* mutant tumor bearing mice shown in panel (A). n=5 per genotype (F) Intra-tumoral pSTAT3 staining by immunohistochemistry for individual tumors upon knockout of *Lif* or *Lifr* in Lkb1-mutant tumors. (G) Schematic representation of LIF neutralization experiment. Autochthonous lung *Lkb1*-mutant tumors were generated by intra-tracheal lentivirus delivery using CRISPR/Cas9 editing in *Kras*^LSL-G12D/+^ *p53*^fl/fl^ Rosa26^LSL-Cas9-P2A-GFP^ mice. Eight weeks after tumor initiation either Anti-LIF neutralizing antibody (700 ug intraperitoneal twice a week) or Isotype control were administered. After three weeks tumor burden was quantified by MRI, lung tissue and bronchoalveolar lavage fluid was collected for analysis. (H) Representative MRI images for mice with *Lkb1*-mutant tumors treated with either Isotype of Anti-LIF neutralizing antibody described in (G). Tumors are outlined in yellow and highlighted in red. (I) Quantification of tumor burden by MRI in (H) after treatment with Anti-LIF Neutralizing antibody or Isotype control described in (G). (J) Protein levels of LIF, IL-6, CSF3, CXCL1, CCL2 were measured in the bronchoalveolar lavage fluid of mice from (G). (K) ExCITE-seq was performed on sorted immune and tumor cells from the lungs of tumor bearing mice described in (B) (n=2 per genotype). UMAP plots of tumor sub-clusters are shown. The proportion of each cluster is displayed on the right. (L) Heatmap showing expression of indicated genes per tumor cluster. (M) UMAP of tumor clusters demonstrating high versus low expression of Inflammatory Response pathways with quantification on the right panel. Results are displayed as mean ± SEM. n of 6 - 9 mice were used for each mouse experiment. For pSTAT3 analysis individual tumors are displayed, otherwise individual mice are shown. Statistical analysis was performed using Mann Whitney U or one-way ANOVA with Tukey’s test where appropriate. * p< 0.05 ** p<0.01 *** p<0.001 **** p< 0.0001.

We then sought to evaluate whether blocking LIF signaling in established *Lkb1*-mutant tumors can reverse the immunosuppressive TIME of these tumors. Eight weeks post-tumor initiation we treated animals bearing *Lkb1*-mutant tumors with LIF neutralizing antibody for three weeks (Figure 4G). Consistent with the genetic KO of *Lif*, LIF neutralization significantly reduced tumor burden (Figure 4H and 4I) demonstrating the therapeutic potential of targeting LIF signaling. Analysis of BAL fluid confirmed decreased levels of LIF after neutralization as well as concurrent reduction in inflammatory cytokines and chemokines such as IL-6, CSF3, CXCL1, and CCL2 (Figure 4J).

In order to understand the mechanism by which autocrine LIF signaling promoted tumorigenesis we performed expanded cellular indexing of transcriptomes and epitopes by sequencing (ExCITE-seq) (Mimitou et al., 2019; Stoeckius et al., 2017). We sorted tumor and immune cells based on GFP and CD45 expression from *Lkb1*-mutant GEMMs with *Lif* KO or *Lifr* KO. Based on the ExCITE-seq analysis we divided the cells into 12 clusters (1 tumor and 11 immune clusters; Figure S7A and S7B). To determine the impact of autocrine LIF signaling in tumors we first focused on transcriptional alterations within the tumor cluster. We found that in both the *Lif* KO and *Lifr* KO condition the expression of key cytokines and chemokines involved in myeloid cell recruitment and function were reduced, including *Cxcl3*, *Cxcl5*, *Cxcl7*, *Csf3*, *Il33*, *Vegfa* (Engblom *et al*., 2016; Propper and Balkwill, 2022) (Figure S7C). Next, in our LUAD GEMM we wanted to evaluate whether autocrine LIF signaling had any impact on tumor heterogeneity. To evaluate this we sub-clustered the tumor cells into 8 clusters (Figure 4K and S7D). This analysis revealed that upon KO of *Lif* or *Lifr*, clusters Sox17 and T7 were greatly reduced (Figure 4I). Differential gene expression suggested that cluster 1 reflects a well-differentiated Alveolar Type 2 cell state based on expression of genes such as *Sftpc,* while clusters Sox17 and T7 are more consistent with dedifferentiated tumors suggested by downregulation of *Nkx2-1* (Figure S7D and S7E) (Yang et al., 2022). Looking at individual subclusters, we observed that the majority of cytokines and chemokines regulated by LIF/LIFR signaling (Figure S7C) were primarily driven by the Sox17 and T7 clusters as demonstrated by increased gene expression of *Cxcl3*, *Cxcl5*, *Vegfa*, and *Csf3* (Figure 4L). Finally, pathway analysis revealed that the dedifferentiated clusters labelled Sox17 and T7 have increased expression of an Inflammatory response signature and that this inflammatory signature is diminished upon ablation of *Lif* or *Lifr* (Figure 4M). Overall, these findings suggest that autocrine LIF signaling in tumors drives the emergence of heterogenous and dedifferentiated tumor subpopulations with high inflammatory signaling.

### Tumor-derived LIF signaling generates an altered TIME rich in immunosuppressive myeloid cells

Given the transcriptional changes in tumor cells and alleviation of inflammatory mediators by ablation of either *Lif* or *Lifr* in *Lkb1*-mutant tumors, we decided to evaluate the impact of LIF signaling on myeloid populations utilizing the ExCITE-seq dataset. To determine the differences in macrophages we clustered macrophages into 9 unique clusters (Figure 5A and S8A). Antibody-derived tags, along with gene expression analysis, allowed us to confidently identify clusters Alveolar MΦ1, cycling MΦ, and ISG^Hi^ MΦ as AMs and CX3CR1^+^ MΦ, Spp1^+^ MΦ, mo- MΦ1, mo-MΦ2, Lyve1+ MΦ, and Arg1^+^ MΦ as IMs based on CD169/CD11c and CD11b/CD14 surface expression respectively (Casanova-Acebes *et al*., 2021; Chakarov et al., 2019; Schyns et al., 2019; Ural et al., 2020) (Figure 5A, S8B, S8C and S8D). In both *Lif* KO and *Lifr* KO conditions we observed an increase in Alveolar MΦ1 and ISG^Hi^ MΦ populations which express high levels of interferon stimulated genes such as *Cxcl9* and *Isg15* (Figure 5A and S8A). These findings are consistent with previous work that showed that LIF signaling leads to downregulation of *Cxcl9* expression in macrophages and that this in turn can promote immune suppression (Pascual-Garcia *et al*., 2019). Furthermore, we saw a dramatic reduction of *Arg1^+^* MΦ and mo-MΦ1 populations and a modest decrease in *Spp1^+^* MΦ and mo-MΦ2 in *Lif* KO and *Lifr* KO conditions (Figure 5A). Notably, *Arg1* expression was significantly reduced in multiple macrophage clusters when LIF signaling was disrupted (Figure 5B), consistent with a reversal of the immunosuppressive state of macrophages induced by *Lkb1*-mutant tumors. To validate our findings from the single-cell studies, we performed immunofluorescent staining and observed a significant decrease in intra-tumoral ARG1 expression in *Lif* KO and *Lifr* KO conditions (Figure 5C and 5D). We next assessed the impact of LIF neutralization on subset of macrophages in *Lkb1*-mutant tumors. Consistent with genetic KO of *Lif* and *Lifr*, LIF neutralization resulted in a significant increase in AMs (Figure 5E). While we did not observe any change in total IM infiltration upon knockout or neutralization of LIF (Figure S9A), intracellular staining of IMs revealed that ARG1 expression was significantly reduced (Figure 5F and 5G), further highlighting the critical role of this cytokine in the modulation of myeloid function.

**Figure 5:**
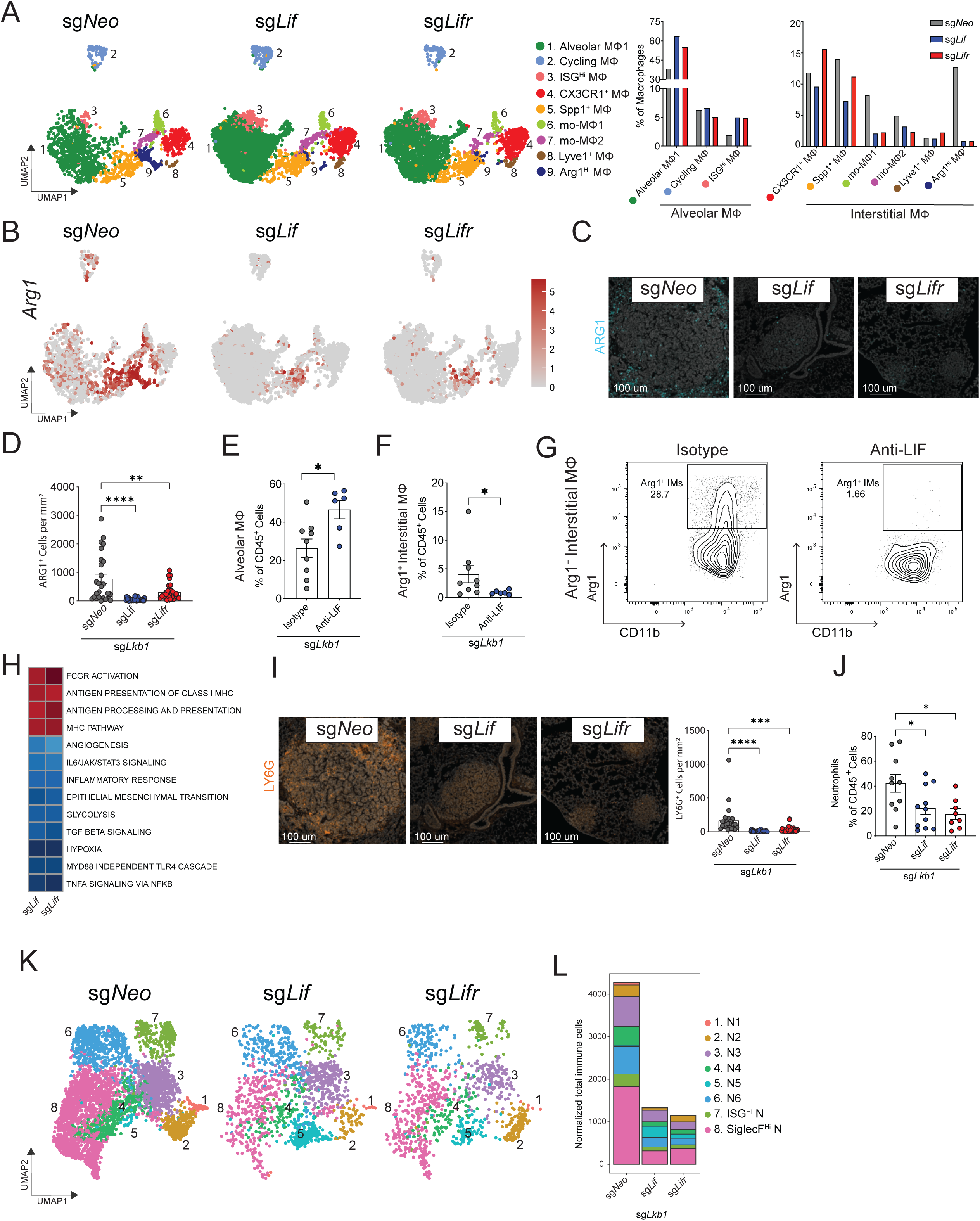
Tumor-derived LIF signaling alters the myeloid composition of the TIME. (A) UMAP plots of macrophage sub-clusters colored by cluster identity in *Lkb1*-mutant tumor bearing lungs comparing control (sg*Neo*) to knockout of *Lif* or *Lifr* (n=2 per condition). The proportion of each cluster, divided into alveolar and interstitial macrophages (Mϕ), is displayed on the right (B) UMAP plots of *Arg1* expression in macrophage subclusters. (C) Immunofluorescence images showing ARG1 (cyan) staining in *Lkb1*-mutant tumors comparing control (sg*Neo*), sg*Lif*, and sg*Lifr*. Scale bar, 100 μm. (D) Quantification of intra-tumoral positive ARG1^+^ normalized per area of images shown in panel (C). (E, F, G) Quantification by flow cytometry of alveolar macrophages (E) and ARG1+ interstitial macrophages (F) as a proportion of noncirculating CD45^+^ cells in the *Lkb1* mutant tumor bearing lungs after LIF neutralization from Figure 4G. (G) Representative flow cytometry plots of ARG1^+^ expression in interstitial macrophages. (H) Significantly upregulated or downregulated pathways (FDR <0.25) in macrophages comparing knockout of *Lif* or *Lifr* to WT from ExCITE-seq data shown in (A). (I) Immunofluorescence images showing Ly6G (orange) staining in *Lkb1*-mutant KP tumors comparing control (sg*Neo*), sg*Lif*, and sg*Lifr*. Scale bar, 100 μm. Quantification of intra-tumoral staining is shown to the right. (J) Quantification by flow cytometry of neutrophils as a proportion of noncirculating CD45^+^ cells in the *Lkb1*-mutant tumor bearing lungs after knockout of *Lif* or *Lifr*. (K) UMAP plots of neutrophil subclusters by ExCITE-seq and colored by cluster identity in *Lkb1*-mutant tumor bearing lungs comparing control (sg*Neo*) to knockout of *Lif* or *Lifr* (n=2 per condition). (L) Quantification of neutrophil clusters seen in (K) by genetic condition. Results are displayed as mean ± SEM. n of 6 - 11 mice were used for flow cytometry data. For Ly6G and ARG1 quantification, individual tumors are displayed. Statistical analysis was performed using Mann Whitney U or one-way ANOVA with Tukey’s test where appropriate. * p< 0.05 ** p<0.01 *** p<0.001 **** p< 0.0001.

Since the above findings suggested that *Lkb1*-mutant tumors altered macrophage function, we further delved into the phenotype and transcriptional heterogeneity of macrophage populations in our tumor models. First, we looked at pathways enriched in Arg1^+^ vs Arg1^-^ macrophages. We found that gene expression in Arg1^+^ macrophages reflected the same pro-inflammatory, immunosuppressive signature that characterized *Lkb1*-mutant tumors (Figure 3A, S9B, and S9C). Consistent with the idea that these transcriptional signaling pathways (IL6/JAK/STAT3 signaling, epithelial mesenchymal transition, hypoxia, TNFA signaling, and TGF beta signaling) are driven by autocrine LIF signaling in tumors, we found that many of these transcriptional changes are reversed with either *Lif* KO or *Lifr* KO (Figure 5H). Ablation of LIF signaling also led to upregulation of transcriptional programs associated with antigen processing and antigen presentation among macrophages isolated from tumor-bearing lungs, further supporting the idea that autocrine LIF signaling promotes tumor immune evasion (Figure 5H).

Next, we characterized neutrophil populations after disruption of LIF signaling. Immunofluorescence and flow cytometric analysis revealed robust reduction in neutrophils in *Lif* KO and *Lifr* KO conditions compared to *Lkb1*-mutant tumors harboring a control gRNA (Figure 5I and 5J). Sub-clustering of neutrophils from our ExCITE-seq analysis revealed a decrease in most neutrophil populations, including the immunosuppressive SiglecF^Hi^ neutrophils in both *Lif* KO and *Lifr* KO tumors (Figure 5K, 5L and S9D). Taken together, these data suggest that autocrine LIF signaling in *Lkb1*- mutant lung tumors drives the development of a pro-inflammatory niche through the transcriptional reprogramming and recruitment of immunosuppressive macrophages and neutrophils.

### LIF signaling suppresses anti-tumor T cell response

Since ablation of LIF signaling diminished the numbers of immunosuppressive myeloid cells within the TIME of *Lkb1*-mutant tumors and led to notable reprogramming of pro-tumorigenic macrophages, we wanted to evaluate whether this led to improved T cell responses in the lungs of these mice. Analysis of T cell populations in the TIME of *Lif* KO and *Lifr* KO tumors demonstrated an increase in IFNγ and TNFα production by CD4^+^ and CD8^+^ T cells from *Lif* KO and *Lifr* KO tumors, consistent with the notion that neutralization of LIF signaling in TIME improves T cell effector function (Figure 6A).

**Figure 6:**
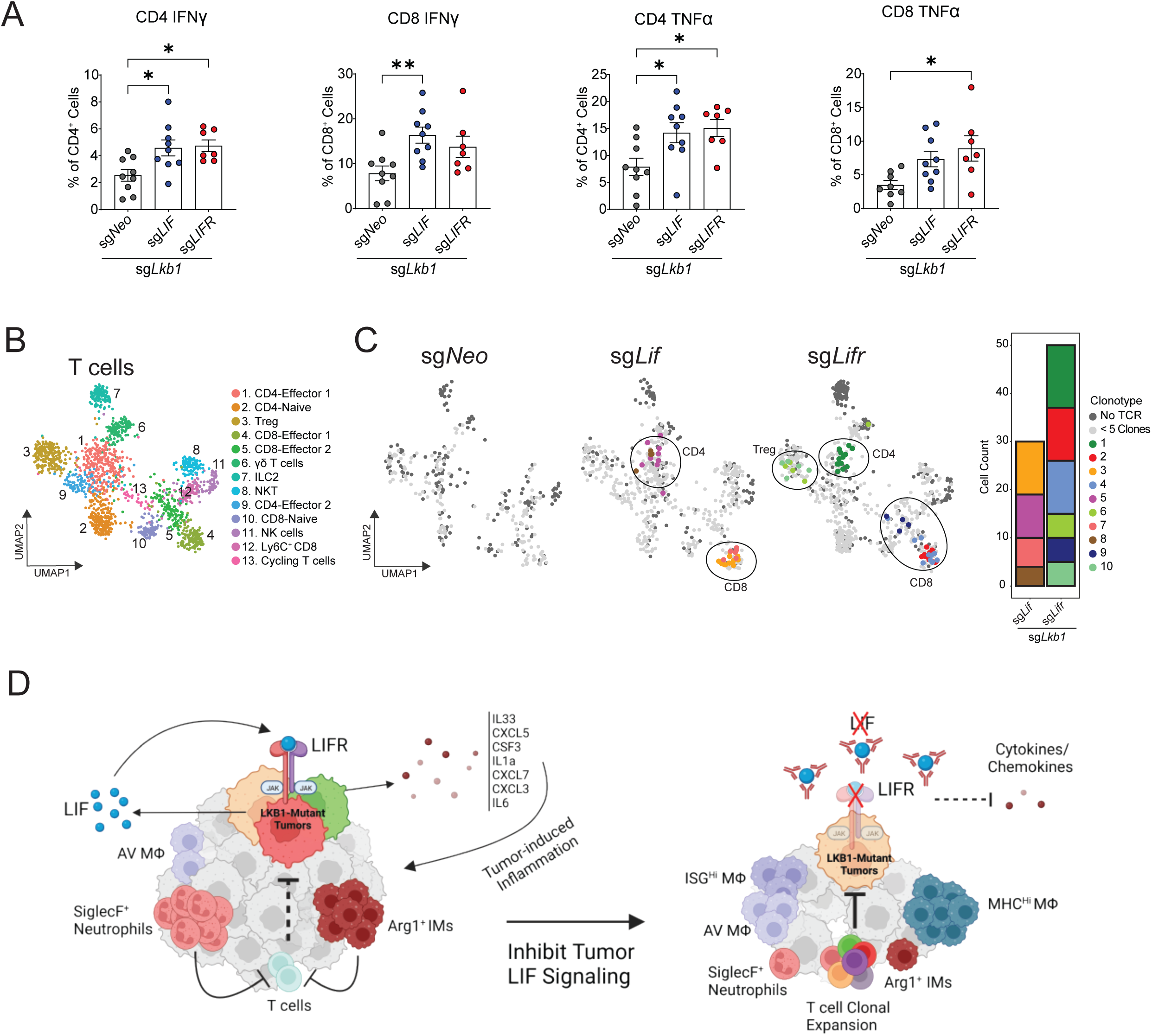
LIF signaling induces an immunosuppressive microenvironment and suppresses T cell responses. (A) T cells were isolated from the lungs of mice with *Kras*^G12D/+^ p53^-/-^ tumors with loss of *Lkb1* and *Lif* or *Lifr* KO, then stimulated with PMA/Ionomycin. Cytokine production of CD4^+^ and CD8^+^ T cells by FACS was plotted for IFNγ and TNFα. (B) UMAP plots of NK and T cell clusters by ExCITE-seq and colored by cluster identity (n=2 mice per condition). (C) UMAP of T cell clusters colored by each clonotype separated by genetic condition: sg*Neo*, sg*Lif*, and sg*Lifr* (n=2). Cells without TCR amplification colored as dark grey and T cells with clonotype numbering < 5 cells colored with light grey. Clonotypes of 5 or more cells are identified with colored dots. Quantification of expanded clonotypes is shown to the right. No expanded clonotypes are seen in the sg*Neo* condition. (D) Schematic of LIF signaling in *Lkb1*-mutant tumors on the tumor immune microenvironment Bar graphs are displayed as mean ± SEM. n of 7 - 9 mice were used for flow cytometry data. Statistical analysis was performed using One-way ANOVA with Tukey’s test. * p<0.05 ** p<0.01 *** p<0.001 **** p< 0.0001.

We then took advantage of the ExCITE-seq dataset to examine the impact of LIF signaling on the adaptive immune system. Clustering of the adaptive immune populations revealed 15 distinct clusters including naïve and effector CD4/CD8 cells, γδ T cells, NK cells, NKT cells, and ILC2s (Figure 6B, S10A). We were able to identify naïve T cells by gene and protein expression of *Lef1*, *Ccr7*, and *CD62L* (Guo et al., 2018), while effector/memory T cells (CD8-Effector 1, CD8-Effector 2, Ly6C+ CD8, CD4-Effector 1, CD4-Effector 2) were identified based on expression of *Id2*, *Nkg7*, *Cxcr3,* and *Ccl5* (Bleul et al., 1997; Cannarile et al., 2006; Hu et al., 2011; Knell et al., 2013; Kurachi et al., 2011; Ng et al., 2020) (Figure S10B). In addition to these genes, we observed distinctive expression of activation/exhaustion markers such as *Pdcd1*, *Tox*, *Tigit,* and *Lag3* in CD8 Effector-1 and CD4 Effector-2 clusters (Figure S10B, S10C, S10D, S10E) (Guo *et al*., 2018). Comparing both *Lif* KO and *Lifr* KO conditions to controls we observed a subtle decrease in Tregs and a reduction in Ly6C^+^ CD8 T cells but otherwise no major changes in T cell clusters in each condition. Given the striking impact of KO of *Lif* or *Lifr* upon tumor progression we speculated that the frequency of clonally expanded antigen-specific T cells increased with ablation of LIF signaling. Analysis of TCR repertoire from *Lif* KO and *Lifr* KO conditions revealed a significant expansion of T cell clonotypes, with clonal expansion primarily restricted to CD8-Effector CD4-Effector, and Treg clusters (Figure 6C and S10F). In contrast, no clonal expansion was observed in *Lkb1*-mutant tumors with intact LIF signaling. Taken together, the observed CD4 and CD8 clonal expansion and the increase in T cell effector function in *Lif* KO and *Lifr* KO tumors suggest that LIF signaling contributes to the suppression of T cell effector responses in *Lkb1*-mutant LUAD.

## Discussion

Human LUAD displays tremendous genetic heterogeneity with tumor mutations impacting prognosis and response to therapy (Arbour *et al*., 2018; Papillon-Cavanagh *et al*., 2020; Ricciuti *et al*., 2020; Shen *et al*., 2019; Wohlhieter *et al*., 2020). ICI, the first line therapy for advanced stage LUAD, impairs tumor progression through induction of anti-tumor immune responses (Gandhi *et al*., 2018). However, ICI are less effective in some genetic subtypes of LUAD, most notably those with *LKB1* or KEAP1 mutations (Papillon-Cavanagh *et al*., 2020; Ricciuti *et al*., 2020; Wohlhieter *et al*., 2020). The connection between genetic mutations of lung cancer and immune evasion remains to be elucidated and studies into this may inform future therapeutic approaches. While the importance of T cell responses in cancer has been well recognized, the impact of myeloid cells on modulating anti-tumor immune responses in LUAD has not been fully understood. Here, we investigate the complex interactions between the TIME and malignant cells in the context of *LKB1*-deficient LUAD. We demonstrated, using patient and mouse samples, that tumor-intrinsic *LKB1* loss-of-function mutations create a pro-inflammatory niche through autocrine LIF signaling. Our study demonstrates that LIF, a cytokine most notably associated with maintenance of pluripotency, promotes the infiltration of immunosuppressive myeloid cells, leading to the suppression of T cell responses and enhanced tumor growth in *LKB1-*mutant tumors (Figure 6D).

Immune profiling of *Lkb1*-deficient tumors revealed a striking increase in IMs, while AMs were significantly reduced. Limited work has been done to investigate the role of diverse lung macrophage subsets in promoting or restricting tumor growth (Casanova-Acebes *et al*., 2021; Fu *et al*., 2022). We found that immunosuppressive markers, including *Arg1*, are predominantly restricted to IMs. Furthermore, we observed that these IM populations are enriched in both GEMMs and human tumors with *LKB1* mutations. Myeloid cells expressing arginase are known to impair both T cell expansion and effector function through the consumption of extracellular arginine (Geiger *et al*., 2016; Miret *et al*., 2019; Rodriguez *et al*., 2004). Our data suggests that *Lkb1*-mutant tumors evade T cell immune surveillance by recruiting immunosuppressive myeloid cells such as Arg1^+^ IMs. To validate the role of *Arg1* in suppressing T cell responses in our *Lkb1*-mutant GEMM further studies are needed using arginase inhibitors that are currently in clinical development (Miret *et al*., 2019; Steggerda et al., 2017).

Our analysis of LUAD patient biospecimens and mouse models demonstrates that *LKB1*-mutant tumors have an elevated number of neutrophils consistent with previously reported results (Koyama *et al*., 2016a). The data presented here further advances our understanding of the consequence of increased neutrophils by the finding that a significant proportion of these cells express *SiglecF*. Prior studies have shown SiglecF^+^ neutrophils to promote tumor growth in orthotopic transplant models of *Lkb1* WT tumors (Engblom *et al*., 2017; Pfirschke *et al*., 2020). Here we demonstrate that *Lkb1*-mutant tumors promote the recruitment of these immunosuppressive myeloid cells into the TIME. While the precise mechanism leading to increased *SiglecF* expression in neutrophils in our mouse model remains to be elucidated, prior work has implicated the tumor-derived CXCR2 ligand CXCL5 in SiglecF*^+^* neutrophil infiltration (Simoncello et al., 2022). Overall, we found that tumor genetics can play a major role in reprogramming of the TIME, by altering the macrophage and neutrophil composition and transcriptional program to create an immunosuppressive microenvironment.

Next, we sought to define the mechanism by which *Lkb1*-mutant tumors promote a pro-inflammatory TIME. Using RNA-seq and multiplex cytokine arrays we demonstrate that *Lkb1* loss leads to upregulation of several chemokines and cytokines. While some of these cytokines have been previously reported to be augmented in *LKB1*-mutant tumors or upon loss of downstream Salt Inducible Kinases (SIK) (Hollstein *et al*., 2019; Koyama *et al*., 2016a), the upregulation of LIF in LKB1-deficient tumors is a novel observation. In agreement with our study, Compton *et al*. demonstrate that LKB1 regulates LIF through SIK/CRTC2 signaling and that inflammatory stimuli can further augment *LIF* expression. The immunomodulatory nature of LIF has been previously shown in autoimmune diseases, embryo implantation and transplantation (Cao et al., 2011; Linker et al., 2008; Stewart et al., 1992b; Wang et al., 2022; Zhang et al., 2019). However, the autocrine role of LIF in cancer and in the regulation of the infiltration and transcriptional state of myeloid cells in the TIME has not been previously described. Using ExCITE-seq we found that the autocrine LIF signaling promotes the expansion of specific tumor sub-clusters with high expression of inflammatory response pathways. This inflammatory signature was primarily driven by two distinct tumor clusters: Sox17 and T7, which were associated with a dedifferentiated state defined by loss of Nkx2-1 expression. This data suggests that LIF signaling may be important in promoting lineage plasticity and the emergence of dedifferentiated tumor populations that drive a pro-inflammatory niche. The role of STAT-driven inflammatory signaling in promoting lineage plasticity has also been observed in other cancer types and requires further investigation (Chan et al., 2022).

Given the profound changes in inflammatory signal upon ablation of LIF signaling, we aimed to characterize the impact of autocrine LIF signaling on the immune infiltration using ExCITE-seq. Our analysis revealed that in particular *Arg1* expression in IM populations is predominantly dependent on autocrine LIF signaling in the tumor. Disruption of LIF signaling in *Lkb1*-mutant tumors not only lead to abrogation of *Arg1* expression in macrophages, but also lead to transcriptional upregulation of genes involved in antigen processing and presentation, which may contribute to augmented T cell responses. While *Lkb1*-mutant lung tumors polarize IMs to express *Arg1* through tumor derived LIF signaling, we have not identified the precise mechanism for *Arg1* upregulation. One potential driver of the altered transcriptional program in IMs is IL-33, which has previously been implicated in polarizing bone marrow-derived macrophages towards an immunosuppressive and anti-inflammatory phenotype (Faas et al., 2021; Taniguchi et al., 2020). Consistent with this, we show that *Il33* expression is downstream of LIFR signaling in *Lkb1-*mutant tumors. In contrast, the overall increase in IM infiltration observed in *Lkb1-*mutant tumors, appeared to be independent of LIF signaling as LIF neutralization and KO of *Lif* or *Lifr,* failed to impact the recruitment of IM and only affected their transcriptional program. Therefore, it is likely that another tumor-derived factor produced by *Lkb1*-mutant tumors is responsible for promoting the infiltration of IMs.

We found that autocrine LIF signaling in tumors regulates the expression of CXCR2 ligands, as well as cytokines including *Csf3* and *Il6,* which are involved in neutrophil recruitment and development (Engblom *et al*., 2016; Forsthuber et al., 2019; Johnson et al., 2018; Mehta et al., 2015). Accumulation of immunosuppressive neutrophils in LUAD is known predictor of poor responses to therapy and poses a major clinical challenge (Hedrick and Malanchi, 2022; Kargl et al., 2017). There are no effective clinically approved therapies to target specifically immunosuppressive neutrophils. Here we demonstrate that inhibition of LIF signaling in tumor cells using either genetic ablation of *Lif* or *Lifr*, or LIF neutralization prevented accumulation of immunosuppressive neutrophils in tumor bearing lungs. These results suggest that blunting LIF signaling pathway may be a promising avenue for targeting this immunosuppressive population of granulocytes in *LKB1*-mutant LUAD.

First, our study highlights that tumor-intrinsic mutations can dictate the inflammatory tone of the immune microenvironment of lung tumors, specifically the pro-tumor polarization of macrophages and neutrophils. By analyzing the transcriptional program of macrophages in *Lkb1*-mutant tumors, we identified an immunosuppressive signature predictive of survival in LUAD patients. Most importantly, we discovered that LIF is major regulator of a pro-inflammatory tumor niche that is responsible for promoting an immunosuppressive TIME of *Lkb1*-mutant tumors. We found that inhibition of LIF reversed some of the immune-evasive characteristics of *Lkb1*-mutant lung tumors as demonstrated by reduction in inflammatory cytokines and chemokines, alteration of the myeloid immune infiltration, improved T cell function, and overall reduced tumor burden, thereby demonstrating that targeting LIF is a viable therapeutic strategy. There is currently an ongoing clinical trial using neutralizing antibodies against LIF in pancreatic cancer (NCT04999969). Results from our study suggest that stratification of patients based on their *LKB1* mutation status is critical for identifying patients that will benefit from this therapeutic approach. Furthermore, given that inhibition of tumor LIF signaling enhanced T cell function and promoted expansion of antigen specific T cells, LIF neutralization may be used to sensitize tumors to ICI, not only in LUAD, but also other tumor types characterized by increased LIF. Additional novel therapeutic strategies to consider in cancers with active LIF signaling include targeting JAK/STAT signaling or *Arg1*. Overall the findings presented here demonstrate the critical role of LIF in tumors as a major regulator of inflammation and a driver of an immunosuppressive TIME and suggest that LIF signaling is a promising therapeutic target for cancers characterized by upregulation of this cytokine.

## Acknowledgements

R. P. was supported by the William Rom fellowship, the Stony Wold-Herbert Fund, and NIH training grant T32 CA009161 and T32 AI100853. T. P. is supported by NIH grants (R37CA222504 and R01CA227649) and an American Cancer Society Research Scholar Grant (RSG-17-200-01–TBE). Work in S.B.K. laboratory was supported by NIH (R01HL-125816), LEO Foundation Grant (LF-OC-20-000351), NYU Cancer Center Pilot grant (P30CA016087). The NYULH Center for Biospecimen Research and Development, Histology, and Immunohistochemistry Laboratory (RRID:SCR_018304), is supported in part by the Laura and Isaac Perlmutter Cancer Center Support Grant; NIH/NCI P30CA016087 and the National Institutes of Health S10 Grants; NIH/ORIP S10OD01058 and S10OD018338. We thank members of the Experimental Pathology Research Laboratory, which is partially supported by the Cancer Center Support Grant P30CA016087 at NYU Langone’s Laura and Isaac Perlmutter Cancer Center. The Akoya Vectra Polaris multispectral scanning system was awarded through the shared instrument grant S10 OD021747.

## Methods

### Mice

All mouse experiments described in this study were approved by the NYU Institutional Animal Care and Use Committee (IACUC). *Kras*^LSL-G12D/+^; *Trp53*^fl/fl^ Rosa26^LSL-Cas9-P2A-GFP/ LSL-Cas9-P2A-GFP^ and Kras^LSL-G12D/+^ *Lkb1*^fl/fl^ (KL) mice have already been described(Koyama et al., 2016b; Lignitto et al., 2019; Platt *et al*., 2014; Sánchez-Rivera *et al*., 2014). For all mouse studies ≥ 5 mouse were used for each experimental condition. Mice with appropriate genotype aged 6 – 10 weeks were randomly selected to begin tumor initiation studies with the USEC lentivirus (Lignitto *et al*., 2019) cloned with paired guides (Vidigal and Ventura, 2015) sg*Neo1*sg*Neo2*, sg*Lkb1*sg*Neo2*, sg*Neo1*sg*Keap1*, sg*Lkb1*sg*Keap1* or sg*Lkb1*sg*Neo*, sg*Lkb1*sg*Lif*, sg*Lkb1*sg*Lifr*. Mice were opened at either 6 weeks or 11 weeks post tumor initiation. Both male and female mice were used equally per experimental arms. (Add Lif neutralization methods)

Prior to sacrifice mice were sedated with ketamine and xylazine. Mice were injected with 2ug of APC anti-CD45 (Biolegend 30-F11) diluted in 100 uL PBS retro-orbitally. The chest of the mouse was opened three minutes after antibody injection. A catheter was inserted into the trachea and 1 mL of saline was injected into the airway. Bronchoalveolar lavage fluid was collected. Fluid was centrifuged at 1500 rpm for 5 minutes and supernatant was collected. Blood was aspirated by cardiac puncture into EDTA tubes. BAL fluid and plasma were stored at -80 C. Lungs were removed and each lobe was separated. Each lobe was cut in half and one set was inflated with 10% zinc formalin for 48 hours, washed in PBS, and resuspended in 70% ethanol prior to embedding. The other half was digested into a single cell suspension first by mincing the tissue on a glass slide followed by digestion with collagenase IV (Sigma Aldirch C5138), DNAse I (Life technologies)

### Anti-LIF antibody generation

A recombinant anti-LIF antibody that cross-reacts with human and mouse LIF was produced from CHO cells using vectors encoding synthetic genes for the heavy chain (Genbank ID: QCA58562.1) and the light chain (Genbank ID: QCA58557.1) from US Patent 10206999-B2 by Biointron. Its binding to LIF was confirmed using recombinant LIF (ACRO Biosystems, LIF-H52H3) on an Octet RED 96e biolayer interferometry instrument (Sartorius).

### Cell lines

KP1233 and 1234 LUAD cell lines were obtained from the laboratory of Tyler Jacks. HY19636 pancreatic cancer cell line was obtained from the laboratory of Alec Kimmelman. Cell lines were mycoplasma tested (PlasmoTest, InvivoGen) and maintained in DMEM (Cellgro, Corning) supplemented with 10% FBS (Sigma Aldrich) and gentamycin (Invitrogen). Cell lines with knockout of *Lkb1*, *Lif*, or *Lifr* were generated by transducing cells with the plasmid lenticrispr V2 puro (Addgene #98290) cloned with a specific guide. Two days after transduction cells were selected with 8 ug/mL puromycin for 5 days.

### Cloning/Virus generation

Cloning of CRISPR sgRNAs was performed as previously described into USEC or PSECC vectors(Sánchez-Rivera *et al*., 2014; Shalem et al., 2014). Lentivirus was generated by co-transfection of HEK293 cells with viral vector and packaging plasmids psPAX2 (Addgene 12260) and pMD2.G (Addgene 12259) using JetPrime transfection reagent (101000046). Media containing virus was collected 72 hours after transfection and filtered through 0.45 uM filter. For *in vivo* experiments the virus was concentrated by ultracentrifugation at 25000 rpm for 2 hours at 4°C. Virus pellet was resuspended in PBS and stored at -80°C until use. Virus was quantified using the GreenGo reporter cell line by adding a serial dilution of the virus directly to cells and measuring percentage of GFP expressing cells at 48 hours. For *in vitro* use, media containing virus was added directly to recipient cells with polybrene (Milipore) 8 ug/mL.

### Flow cytometry

Single cell suspensions were initially stained with UV zombie fixable viability dye (Biolegend 423107) for 15 minutes at room temperature per manufacturer’s protocol. Cells were then resuspended in FACS buffer (PBS 0.5% BSA, 1 mM EDTA, 0.1% Sodium Azide) and incubated with Fc block (2.4G2, Bioexcell) for 10 minutes at 4°C. Cells were then incubated with surface antibodies at 15 minutes or 30 minutes at 4°C depending on the antigen. To stain for transcription factors, cells were permeabilized and fixed using the FoxP3 Staining buffer kit (eBioscience 00552300). Intracellular staining was performed by blocking with Fc for 10 minutes followed by 30 minutes of antibody staining at room temperature.

For intracellular cytokine staining single cell suspensions were stimulated with PMA (0.1 ug/mL, Sigma P-8139), ionomycin (1ug/mL, Sigma I-0634), Golgi Plug (BD Biosciences 55029, 1:1000), and Golgi Stop (BD biosciences 555029, 1:1000) for 3.5 hours in RPMI with 10% FBS at 37°C. Cells were washed and stained for surface markers as described above. For intracellular staining cells were fixed initially with 2% PFA for 10 minutes at room temperature and then permeabilized with 0.5% saponin for 15 minutes at room temperature. Cells were than incubated in 0.5% saponin with Fc block for 10 minutes and then intracellular antibody staining for 30 minutes at room temperature. Cells were then resuspended in FACS buffer. Samples were analyzed on the BD LSR Fortessa Cell Analyzer.

### qPCR

RNA was collected from tumor cell lines using the RNeasy mini kit (Qiagen). cDNA was synthesized from mRNA using SuperScript VILO (Invitrogen) per manufacturers protocol. qPCR was performed using the SybrGreen master mix (Applied biosystems).

### *Lifr* Knockout Validation by Western blot

For validation of *Lifr* knockout, 1234 KP LUAD cells were plated in a 6 well dish. The next day media was aspirated and replaced with serum free DMEM. After 6 hours of serum starvation cells were stimulated with recombinant mouse LIF (5 ng/mL, Peprotech). Cells were lysed with Pierce RIPA buffer (Thermo Scientific) on ice. Samples were scraped and collected into a microcentrifuge tube. Samples were centrifuged at 10000 rpm at 4°C for 15 minutes. Supernatant was collected and protein was quantified using DC Rad Protein Assay kit. Protein was diluted to 2 ug/uL with water and 4x NuPage LDS sample buffer. Samples were boiled at 95°C for 10 minutes. Proetin was loaded onto Invitrogen 4-12% Bis-Tris gel. Gel was run at 140V for 90 minutes. Transfer was performed on to a nitrocellulose membrane at 100V for 120 minutes. Membrane was blocked using 5% BSA (in TBST) for 60 minutes at room temperature and then incubated with the primary antibodies pSTAT3 (CST 9145, 1:1000) and GAPDH (Santa Cruz 25778, 1:4000) in 5% BSA overnight at 4°C. The membrane was then washed in TBST and incubated with the secondary antibody. To detect the band enhanced chemiluminescent horseradish peroxidase substrate (Thermo Scientific Super Signal West PICO Plus) was added to the membrane for 5 minutes. The membrane was visualized using the General Electric Amersham Imager 680.

### Bulk RNA-seq

Tumor cells were sorted as singlets, live^+^/dead^-^ CD45^-^GFP^+^. RNA was isolated using Purelink RNA mini kit (Invitrogen) per manufacturer’s instruction. cDNA was synthesized from RNA using SMARTer PCR cDNA synthesis kit (Clontech) per manufacturer’s instruction. Sequencing libraries were prepared using Nextera XT DNA library preparation kit (Illumina) per manufacturer’s instruction. Samples were pooled at equimolar ratios. Libraries were loaded on an SP11 cycle flow cells and sequenced on Illumina NovaSeq6000. Read qualities were evaluated using FASTQC (Babraham Institute) and mapping to GRCm38 (GENCODE M25) reference genome using STAR program(Dobin et al., 2013) with default parameters. Read counts, TPM and FPKM were calculated using RSEM program(Li and Dewey, 2011). Identification of differentially expressed genes (DEGs) between different genotype of KP tumor was performed using DESeq2 in R/Bioconductor. All plots were generated using customized R scripts. Hallmark pathways were downloaded from MSigDB(Liberzon et al., 2011). Pathway enrichment analysis was performed using GSEA preranked program(Subramanian et al., 2005) based on log2FC values of all genes.

### Single cell RNA seq/ExCITE seq

Approximately 12000 Lung immune cells from each condition (2 mice per condition) were sorted as live^+^/dead^-^ CD45-circulating^-^ CD45^+^. For ExCITE-seq tumor cells were sorted as live^+^/dead^-^ CD45-circulating^-^ CD45^-^ GFP^+^ and added to immune cells prior to multiplexing. Then samples were multiplexed using cell hashing antibodies. Cells from each sample were pooled and loaded into 10X Chromium. Gene expression together with Hashtag oligo (HTO) libraries were processed using Cell Ranger (v5.0.0) in multi mode. Cell-containing droplets were selected using the default filtering from Cell Ranger count “filtered_feature_bc_matrix”. UMI count matrices from each modality were imported into the same Seurat(Hao et al., 2021; Stuart et al., 2019) object as separate assays. Viable cells were filtered based on having more than 200 genes detected and less than 15% of total UMIs stemming from mitochondrial transcripts. HTO counts were normalized using centered log ratio transformation before hashed samples were demultiplexing using the Seurat::HTODemux function. RNA counts were normalized using Seurat::SCTransform function with regressions of cell cycle score, ribosomal and mitochondrial percentages.

Cells from multiple conditions were combined using Seurat standard scRNSeq integration workflow with 3000 anchor genes. A shared nearest neighbor graph was then built based on the first 40 principal components (PCs) followed by identification of cell clusters using Leiden algoithm and Seurat::FindClusters function at multiple resolutions in order to identify potential rare cell types. Cell types were annotated based on canonical cell type markers and differential expressed genes of each cluster identified using Seurat::FindAllMarkers function with a logistic regression model. Clusters expressing markers of the same cell type were merged into a single cluster. Cell were then projected on to a uniform manifold(McInnes, 2018) using the top 40 PCs for visualization.

Processing of ExCITESeq data was similar to scRNASeq data as described above except 10% of total mitochondrial transcripts was used for cell filtering. Protein counts were normalized using centered log ratio transformation. Multimodal integration was performed using the weighted-nearest neighbor (WNN) method in Seurat. Briefly, a WNN network was constructed based on modality weights estimated for each cell using Seurat::FindMultiModalNeighbors function with top 40 and top 30 PCs from normalized RNA and protein counts, respectively. Differential expression anlaysis for all genes was performed using Mast program and Seurat::FindMarkers function. Pathway enrichment analysis was performed using GSEA preranked program(Subramanian *et al*., 2005) based on log2FC values of all genes.

### Single-nuclei RNA-seq of lung tumor from patients with NSCLC

Nuclei were prepared for 10x Genomics-based single nuclei RNA sequencing analysis according to a previously published protocol(Drokhlyansky et al., 2020) .Briefly, each frozen sample was thawed and macerated in CST buffer for 10 minutes, filtered (70 micron pluriStrainer) and spun at 500g for 5 min at 4C to pellet nuclei. Nuclei were resuspended in the same buffer without detergent, filtered (10 micron pluriStrainer) and counted using AOPI on a Nexcelom Cellometer. Approximately 10,000 nuclei were loaded immediately into each channel of a 10x Chromium chip (10x Genomics) using 5’ V1.1 chemistry according to the manufacturer’s protocol. The resulting cDNA and indexed libraries were checked for quality on an Agilent 4200 TapeStation and then quantified and pooled for sequencing on an Illumina NextSeq 550.

### Immunohistochemistry

Tissues were fixed in 10% zinc formalin for 48 hours and processed through graded ethanols, xylene and into paraffin in a Leica Peloris automated processor. Five-micron paraffin-embedded sections were either stained with hematoxylin and eosin or immunostained on a Leica BondRX® autostainer, according to the manufacturers’ instructions. In brief, sections stained first underwent epitope retrieval for 20 minutes at 100° with Leica Biosystems ER1 solution (pH 6.0, AR9961) followed by a 1 hour incubation with an anti-Lkb1 (1:5000, CST, 13031) or anti-NQO1 primary antibody (Nqo1 (1:100, HPA007308, Sigma-Aldrich) in Leica diluent (Leica, Cat ARD1001EA) and subsequent detection using the BOND Polymer Refine Detection System (Leica, Cat DS9800). All antibody incubations were performed at room temperature. Sections were counter-stained with either hematoxylin and scanned on either a Leica AT2 or Hamamatsu Nanozoomer HT whole slide scanner. Slides were analyzed using QuPath 0.2.3.

### Immunofluorescence

Tissue sections were processed as above. The iterative multiplex immunostaining protocol was performed on the Leica BondRX automated stainer, according to manufacturers’ instructions with the antibodies (see Antibody Table). Briefly, all slides underwent sequential heat retrieval with either Leica Biosystems epitope retrieval 1 solution (ER1, pH 6.0, AR9961) or retrieval 2 solution (ER2, pH 9.0, AR9640), followed by primary and secondary antibody incubations and tyramide signal amplification (TSA) with Opal® fluorophores as shown in Antibody Table. Primary and secondary antibodies were removed during heat retrieval steps while fluorophores remained covalently attached to the epitope. Semi-automated image acquisition was performed on a Vectra® Polaris multispectral imaging system at 20X. Whole slide unmixed scans were viewed with Akoya Phenochart. Slides were analyzed using QuPath 0.2.3.

### Human Immunohistochemistry

Chromogenic immunohistochemistry (IHC) on the human tumor microarray was performed using the following antibodies: unconjugated polyclonal rabbit anti-human myeloperoxidase (IVD, Cell Marque Cat# 289A-78, Lot# 10 unconjugated, RRID: AB_2335990) and rabbit anti-human pSTAT3 clone D3A7 (Cell Signaling Technologies Cat# 9145, Lot# 43 RRID: AB_2491009) raised against a synthetic phosphopeptide corresponding to residues surrounding Tyrosine 705 of murine STAT3. IHC was performed on a Ventana Medical Systems Discovery Ultra platform using Ventana’s reagents and detection kits unless otherwise noted. In brief, five-micron tissue sections were collected onto Plus slides (Fisher Scientific, Cat # 22-042-924), air-dried and stored at room temperature prior to use. Myeloperoxidase was assayed using validated in vitro diagnostic method according to manufacturer’s instructions. Sections for pSTAT3 were deparaffinized online, followed by antigen retrieval in CC1 (TRIS-Borate-EDTA, ph 8.5, Cat# 950-500) for 20 minutes at 99°C. Antibody was diluted 1:100 and incubated for 12 hours followed by detection with goat anti-rabbit Horseradish Peroxidase conjugated multimer (Ventana Medical Systems, Cat# 760-4311) incubated for 8 minutes, and detected with ChromoMap RUO (Cat# 760-159) DAB detection. Slides where washed in distilled water, counterstained with hematoxylin, dehydrated thru graded alcohols, cleared in xylene and mounted with synthetic permanent media. Appropriate positive and negative controls were included with the study sections.

### RNA scope

Tissues were processed as above, and five um sections were cut within 2 days of performing the assays. *In situ* hybridization staining with the LIF probe (ACDBio, 322700) was performed according to the ACDBio protocol (document UM322700) using RNAscope 2.5 LSx Reagent Kit – BROWN (ACDBio, 322700). The slides were counterstained with hematoxylin, coverslipped and scanned on Leica AT2 whole slide scanner at 40X.

### ELISAs

Cytokine/Chemokine multiplex assays were performed on BAL fluid by Eve Technologies using their Mouse Cytokine/Chemokine 31-plex Discovery Assay Array (MD31).

### Human patients’ survival analysis

Clinical and genomic data from the study by Samstein et al. were downloaded from https://www.cbioportal.org/. This cohort included 1,661 patients who had received at least one dose of an ICI (targeting PD-1, PD-L1 or CTLA-4) and who had tumor genomic profiling using the commercially available MSK-IMPACT assay.

Miao et al.: This was a cohort of 249 ICI-treated patients with microsatellite-stable (MSS) solid tumors. Pre-treatment samples were analyzed with whole-exome sequencing (WES). Clinical and genomic data were downloaded from https://www.cbioportal.org/.

### TCGA analysis

RNA-seq gene expression profiles of primary tumors and relevant clinical data of 515 LUAD patients were obtained from The Cancer Genome Atlas(Cancer Genome Atlas Research, 2014) (TCGA, gdac.broadinstitute.org). *STK11 (Lkb1)* mutational status of TCGA tumor samples was retrieved from cBioPortal(Cerami et al., 2012) using the TCGA PanCancer Atlas collection (gdc.cancer.gov/about-data/publications/pancanatlas). Within this dataset of 515 samples, 510 were assigned mutational status as follows: 437 *STK11* WT; 73 *STK11* mutant (missense, splice, or truncating mutations). Patients were grouped by mutational status, as described in the figure legend, and the distribution of standardized LIF expression across groups was illustrated using an Empirical Cumulative Distribution Function plot (ECDF) where significance was assessed using the Kolmogorov-Smirnov test. For survival analyses, patients were stratified based on LIF expression and Kaplan-Meier 5-year survival analyses were conducted to compare high-LIF expressing patients (top 10%, n=51 patients) with the rest of the cohort (n=464 patients), and significance was assessed using the log-rank test. All survival analyses were conducted using the survival package in R. All statistical analyses were conducted in the R statistical programming language (R-project.org).

### Statistical analysis

GraphPad Prism 9 was used for statistical analyses. Data is plotted as mean +/- SEM and a p value of <0.05 was considered significant. Outliers were identified using the Grubb’s method. Experiments with more than 2 experimental arms were analyzed with one-way ANOVA and Tukey’s test for multiple comparisons. Experiments with two arms were analyzed with Mann-Whitney U test with Two-tailed analysis. * - p<0.05, ** - p<0.01, *** - p<0.001, **** p<0.0001.

---

**Figure S1:**
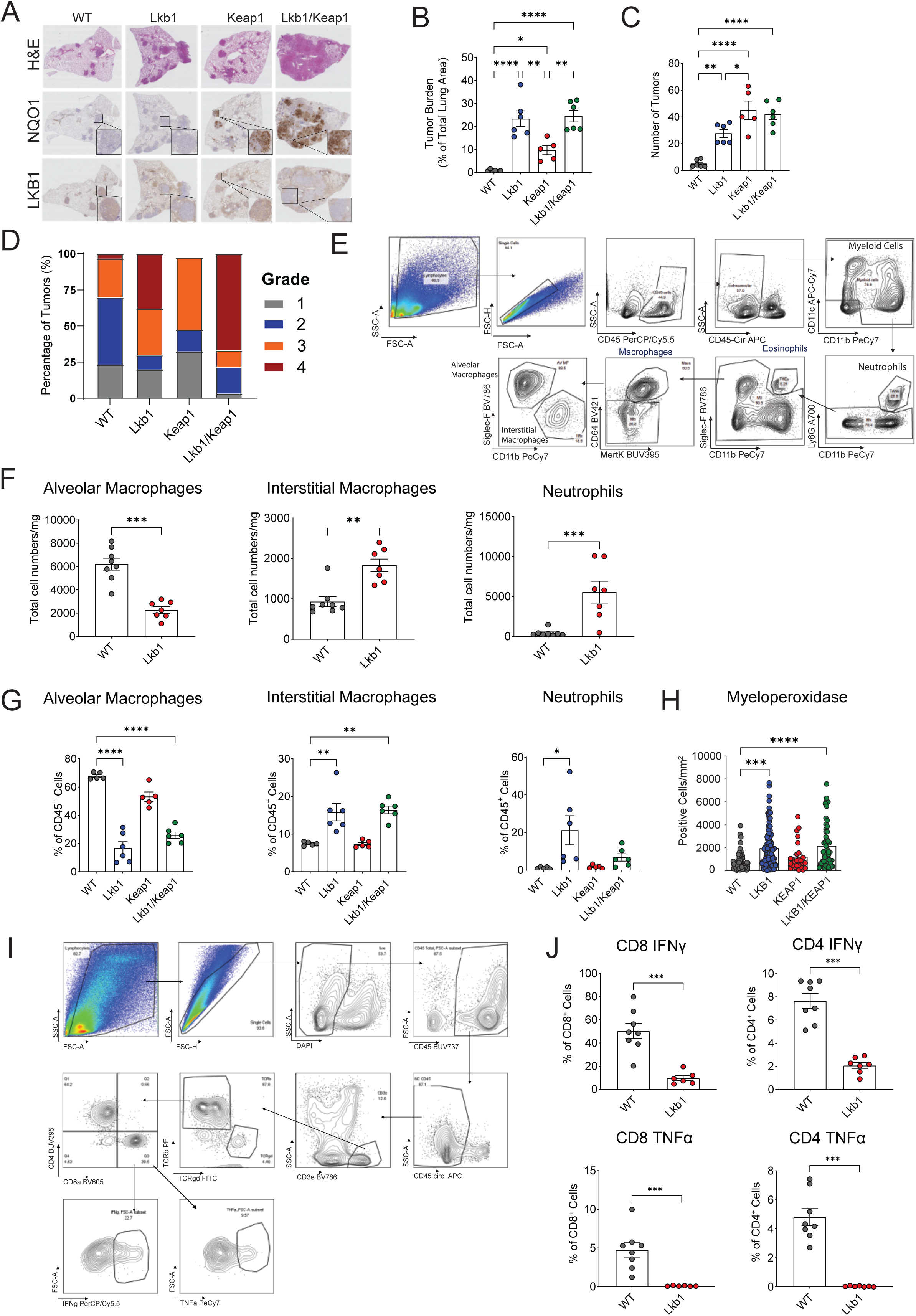
*Lkb1-* and *Keap1*-mutant tumors alter the tumor microenvironment. (A) Images of lung tumor by H&E (top) and NQO1 (middle) and LKB1 (bottom) staining by immunohistochemistry validating *in vivo* CRISPR/Cas9 editing in our genetically engineered *Kras*^G12D/+^ *p53*^-/-^ lung cancer mouse model. Genotypes are indicated. (B) Tumor burden represented as cross-sectional area as a percentage of total lung area measured on a midline H&E section at 11 weeks post tumor initiation (n=5-7). (C) Quantification of tumor number in indicated genotype. n=5-6 per genotype. (D) Percentage of tumors with corresponding grade stratified by genotype. (E) Gating strategy for identification of myeloid cells in tumor bearing lungs. CD45-Cir reflects immune cells labelled by intravascular APC anti-CD45. Myeloid cells were gated as singlets, CD45^+^/CD45-Cir^-^/CD11b^+^/CD11c^+^; Neutrophils were gated as singlets, CD45^+^/CD45-Cir^-^/CD11b^+^/Ly6G^+^; after removing neutrophils and eosinophils, alveolar macrophages were gated on CD64^+^/MertK^+^/SiglecF^+^/CD11b^-^ and Interstitial macrophages were gated on CD64^+^/MertK^+^/SiglecF^-^/CD11b^+^. (F) Total cell numbers of myeloid cells (alveolar macrophages, interstitial macrophages, and neutrophils) in WT and LKB1 mutant lung tumors normalized for lung weight. (n=6-8 per genotype). (G) Myeloid cell infiltration (alveolar macrophages, interstitial macrophages, and neutrophils) represented as a percentage of total tissue infiltrating immune cells (CD45^+^ Vascular CD45-Cir^-^) in each indicated genotype. (H) Quantification of myeloperoxidase (MPO) staining in a human tumor microarray. Individual mice or human tumor cores are shown for all bar plots with mean and standard error. (I) Gating strategy for identification of T cells in tumor bearing lungs. T cells were gated as singlets, live (DAPI^-^), CD45^+^, vascular CD45^-^, CD3e^+^. (J) IFNγ and TNFα production by CD4^+^ and CD8^+^ T cells isolated from *Lkb1*-mutant or wildtype tumor bearing lungs after PMA/Ionomycin stimulation by flow cytometry plotted as a percentage of CD4^+^ and CD8^+^ T cells. n=6-8 per genotype Individual mice or tumors are shown in the bar plots with mean and standard error. Statistical analysis was performed using one-way ANOVA and Tukey’s test or Mann-Whitney U test where appropriate. * p< 0.05 ** p<0.01 *** p<0.001 **** p< 0.0001

**Figure S2:**
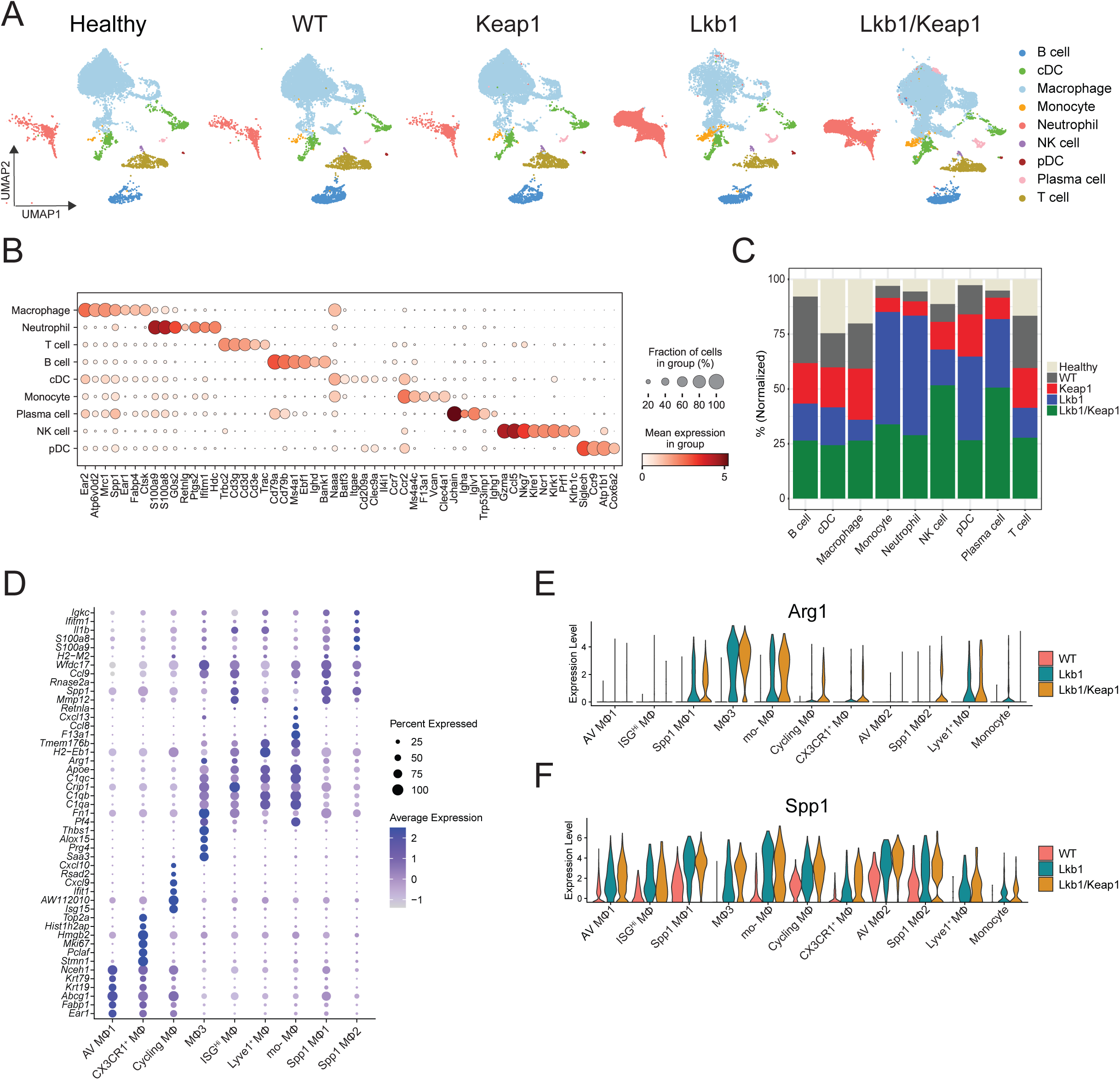
scRNA-seq of immune populations in *Lkb1-* and *Keap1-*mutant lung tumors. (A) UMAP visualization of all immune cells isolated from lungs of healthy mice or mice with tumors from our GEMM LUAD model using single cell RNA-seq. Conditions are stratified by genetic condition. Each colored cluster represents a cell type identified based on gene expression. (B) Top DEGs of immune cells in panel (A). (C) Proportion of immune populations attributed each tumor genotype divided by cell type. (D) Differentially expressed genes of macrophage subclusters. (E, F) Violin plot showing (E) *Arg1* and (F) *Spp1* gene expression in each macrophage sub-cluster among different tumor genotypes.

**Figure S3:**
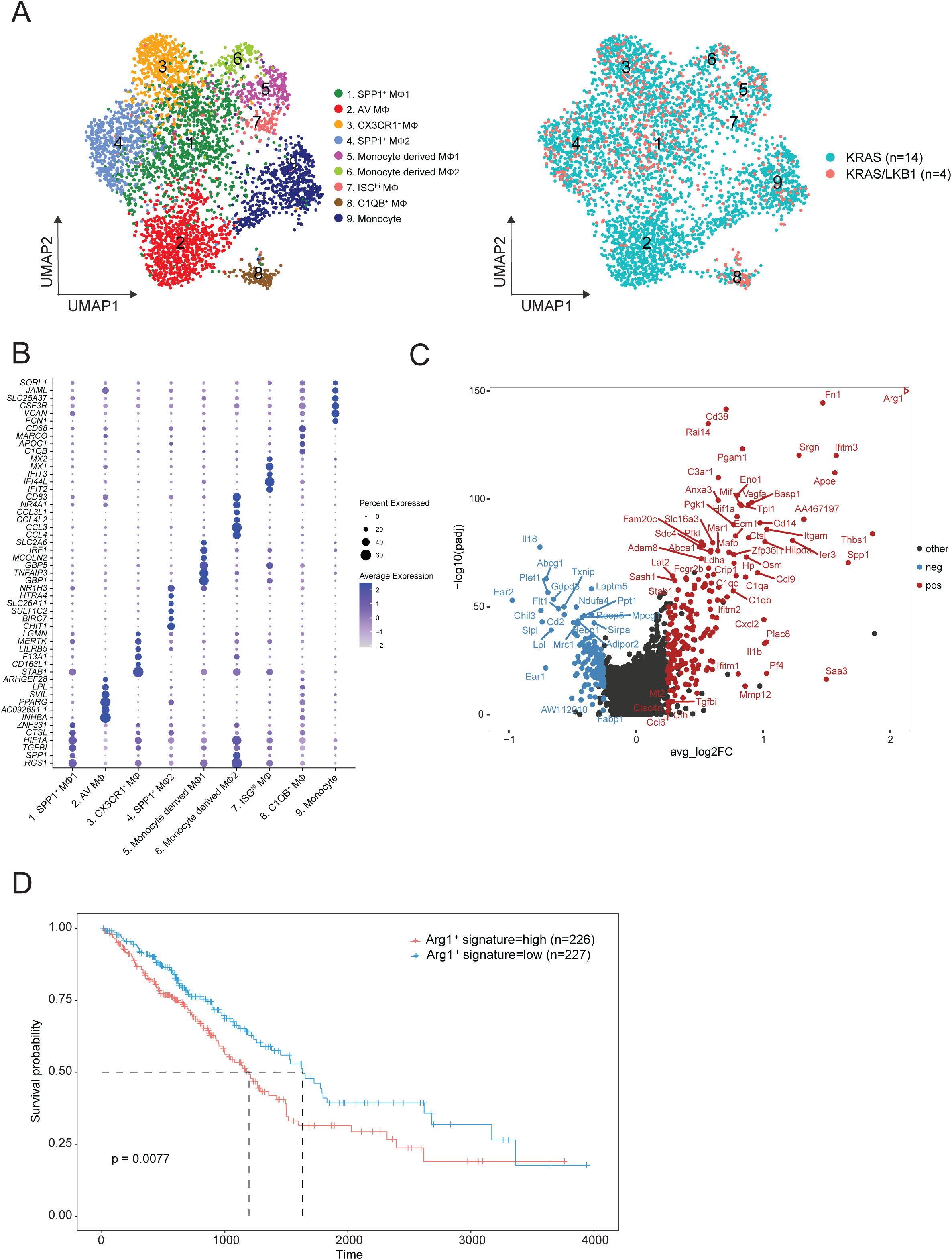
*LKB1*-mutant human tumors alter the infiltration and transcriptional program of myeloid cells. (A) UMAP plot on the left demonstrates visualization of myeloid cell sub-clusters by single cell nuclei RNA-seq of immune cells from human lung tumors. Each colored cluster represents a cell type identified based on gene expression. UMAP plot on the right shows myeloid cell clustering labeled according to tumor genotype. (B) Top differentially expressed genes of myeloid subclusters in panel (A). (C) Volcano plot showing differential gene expression between *Arg1^+^* vs Arg1^-^ macrophages in *Lkb1* mutant condition. Up-regulated genes are highlighted in red and down-regulated genes highlighted blue. Statistical analysis is outlined in Materials and Methods. (D) Survival (Kaplan-Meier) plots of lung adenocarcinoma patients from TCGA with bulk RNA-seq data. Patients were stratified based on high versus low expression of Arg1^+^ macrophages signature generated from (C).

**Figure S4:**
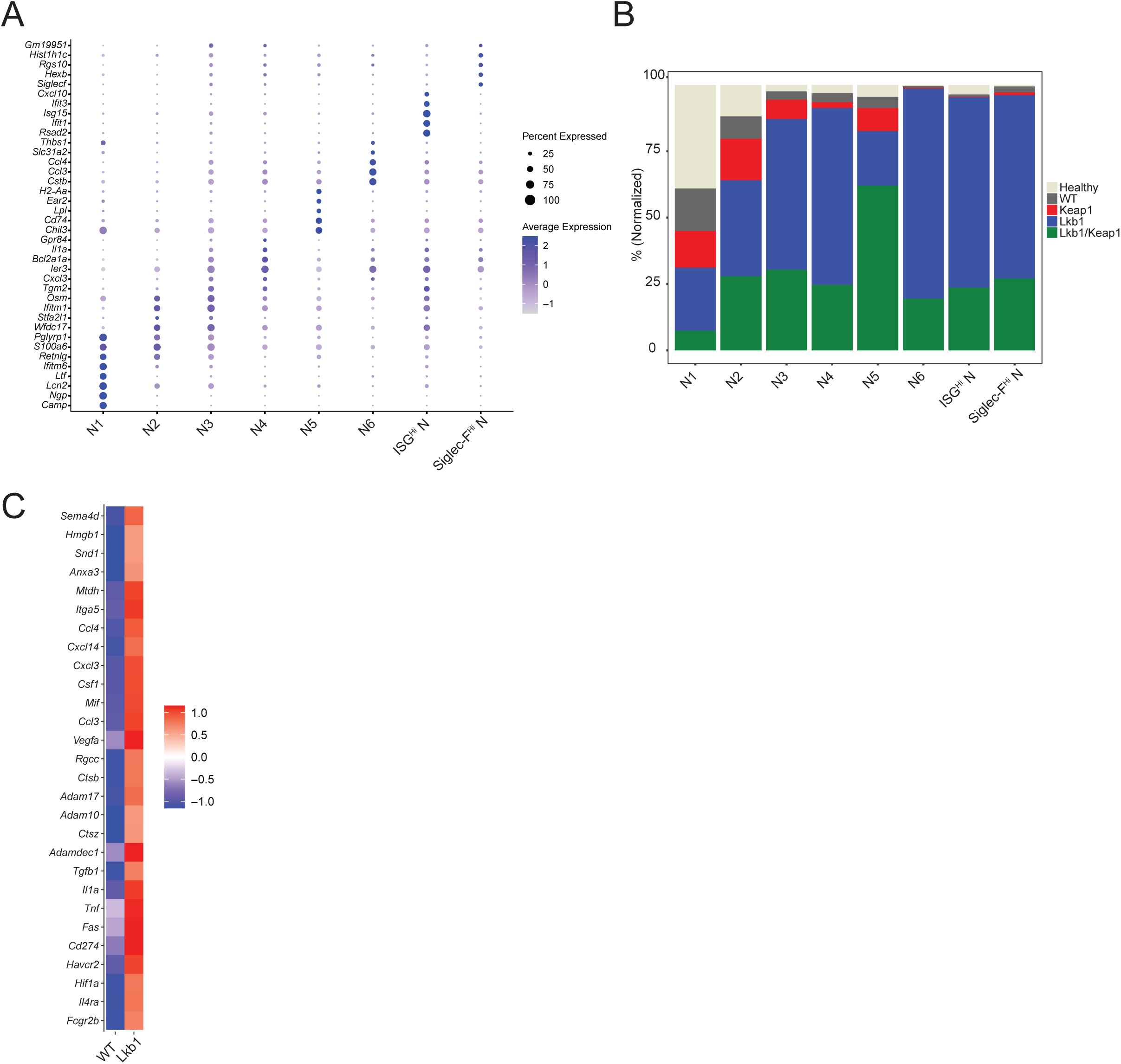
*Lkb1*-mutant tumors have increased neutrophil infiltration with upregulation of immunosuppressive transcriptional program. (A) Top differentially expressed genes of neutrophil subclusters in mouse tumors. (B) Proportion of neutrophil subclusters normalized by total cells for each cluster and divided by tumor genotype. (C) Heatmap of gene expression of selected genes associated with immunosuppression comparing neutrophils from *Lkb1-*mutant to WT lung tumors.

**Figure S5:**
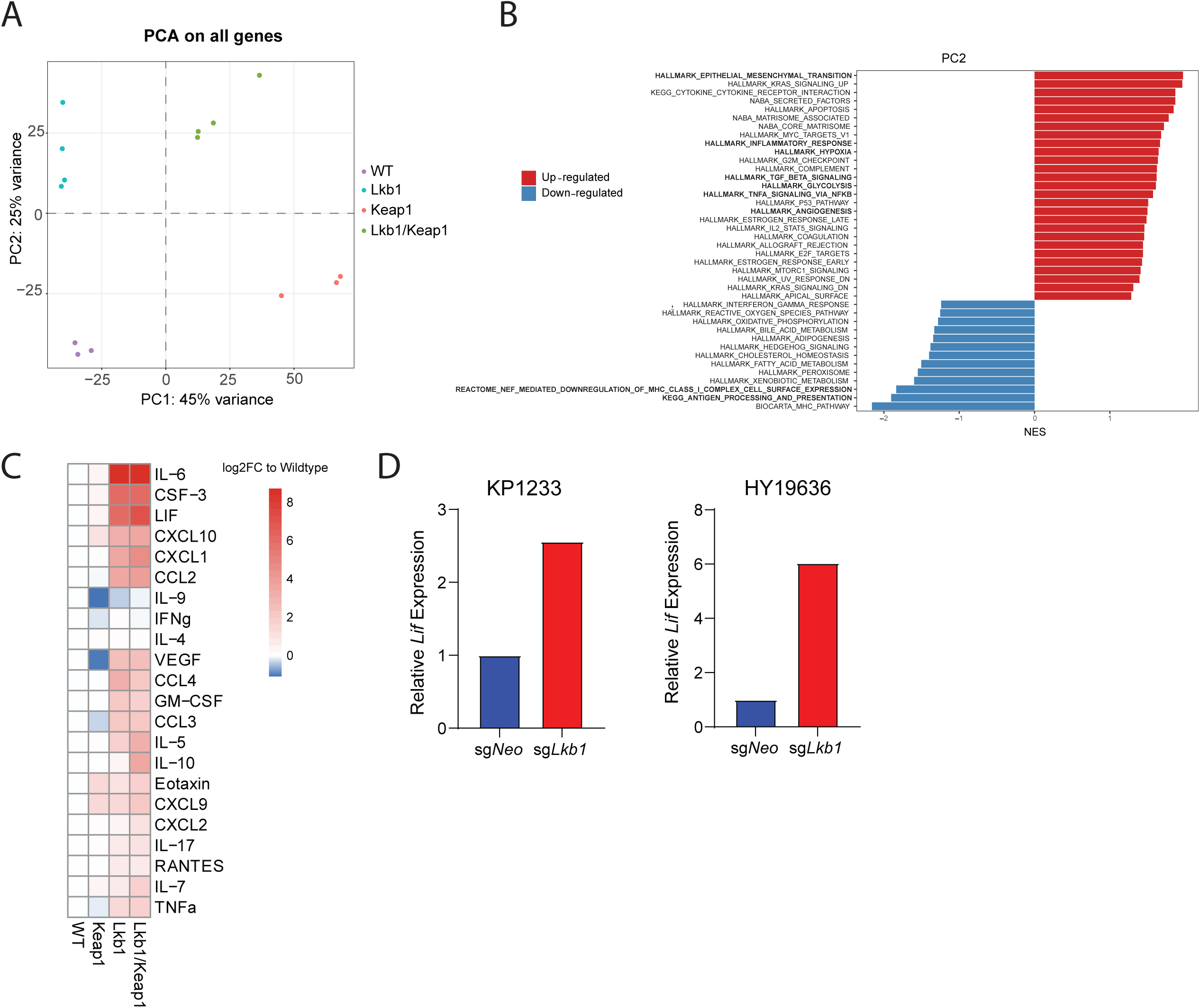
*Lkb1*-mutant tumors have increased inflammatory pathways and cytokine/chemokine levels. (A) RNA-seq was performed on sorted GFP^+^ WT, Lkb1, Keap1, and Lkb1/Keap1 tumor cells. Principal component analysis was performed and PC1 vs PC2 was plotted. (B) GSEA was performed, and top upregulated and downregulated hallmark pathways were plotted for PC2. (C) Chemokine/cytokine multiplex analysis was performed on the bronchoalveolar lavage fluid of tumor bearing mice and log2 fold change of protein levels was plotted relative to WT condition. (D) *Lif* expression by qPCR was measured in *Lkb1-*mutant (*sgLkb1*) and control (sgNeo) KP1233 (lung adenocarcinoma) and HY19636 (pancreatic adenocarcinoma) cell lines.

**Figure S6:**
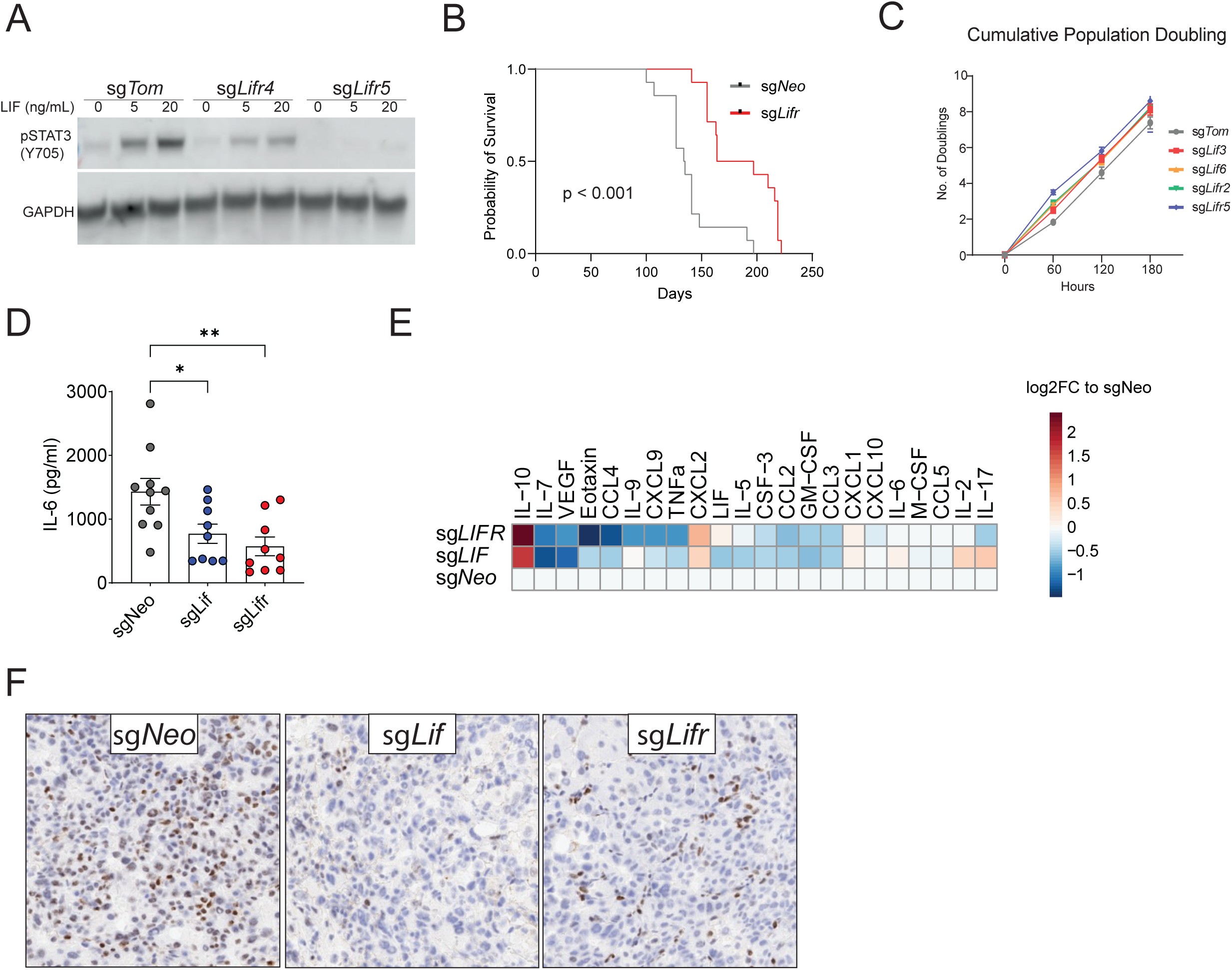
*Lif* or *Lifr* KO reverses immunosuppressive environment of *Lkb1*-mutant tumors. (A) Guides targeting the mouse *Lifr* were validated by stimulating KP mouse 1234 cell lines transduced with control (sgTom), sg*Lifr4*, or sg*Lifr5* vector expressing Cas9 with recombinant mouse LIF for 15 minutes and measuring pSTAT3 by western blot. The sgRNA sg*Lifr*5 was used for subsequent experiments designated as sg*Lifr*. (B) Survival of *Kras*^LSL-G12D/+^ mice induced with tumors with *Lkb1*/*Lifr* KO or control (*Lkb1* KO). (C) Cumulative population doublings of KP mouse cell lines *in vitro* with knockout of *Lif* or *Lifr*. Survival analysis was performed using the Log-Rank (Mantel-Cox) test. (D) IL-6 concentration in bronchoalveolar lavage fluid of *Lkb1* mutant tumor bearing mice. n=9-10 per genotype. (E) Chemokine/cytokine multiplex analysis was performed on the BAL fluid of tumor bearing mice and log2 fold change of protein levels was plotted relative to sg*Neo* condition. (F) Representative intra-tumoral pSTAT3 staining for indicated genotypes (sg*Lkb1/sgLif, sgLkb1/sgLifr* and *sgLkb1/sgNeo*). Individual mice or tumors are shown in the bar plots with mean and standard error. Statistical analysis was performed using one-way ANOVA and Tukey’s test where appropriate. * p< 0.05 ** p<0.01 *** p<0.001 **** p< 0.0001

**Figure S7:**
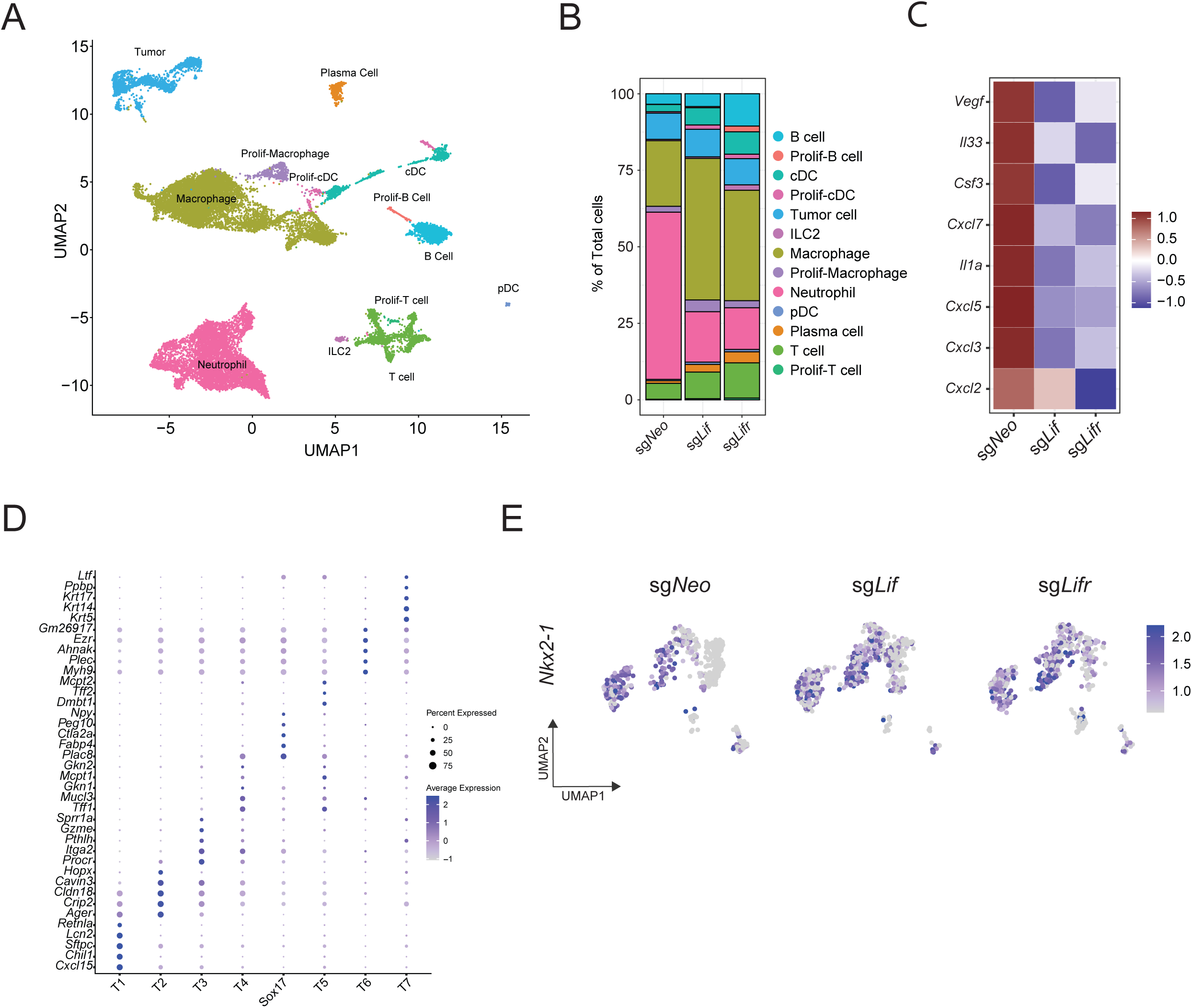
ExCITE-seq reveals disruption of LIF/LIFR-STAT3 signaling alters tumor heterogeneity. (A) ExCITE-seq was performed on sorted immune and tumor cells from our tumor mouse model. UMAP plots of immune and tumor cells is shown. Each colored cluster represent a cell type and identified based on gene expression. (B) Quantification of clusters seen in panel A from *Lkb1*-mutant tumors with *Lif* KO (sg*Lif*), *Lifr* KO (sg*Lifr*) and control (sg*Neo*). (C) Gene expression heatmap of selected genes (*Vegfa, Il33, Csf3, Cxcl7, Cxcl5, Cxcl3, and Cxcl2*) from ExCITE-seq tumor cluster. (D) Differentially expressed genes of tumor subclusters from ExCITE-seq dataset. (E) UMAP plots of *NKx2-1* expression in *Lbk1*-mutant tumor clusters divided by control (sgNeo), *Lif* KO (sg*Lif*), or *Lifr* KO (sg*Lifr*).

**Figure S8:**
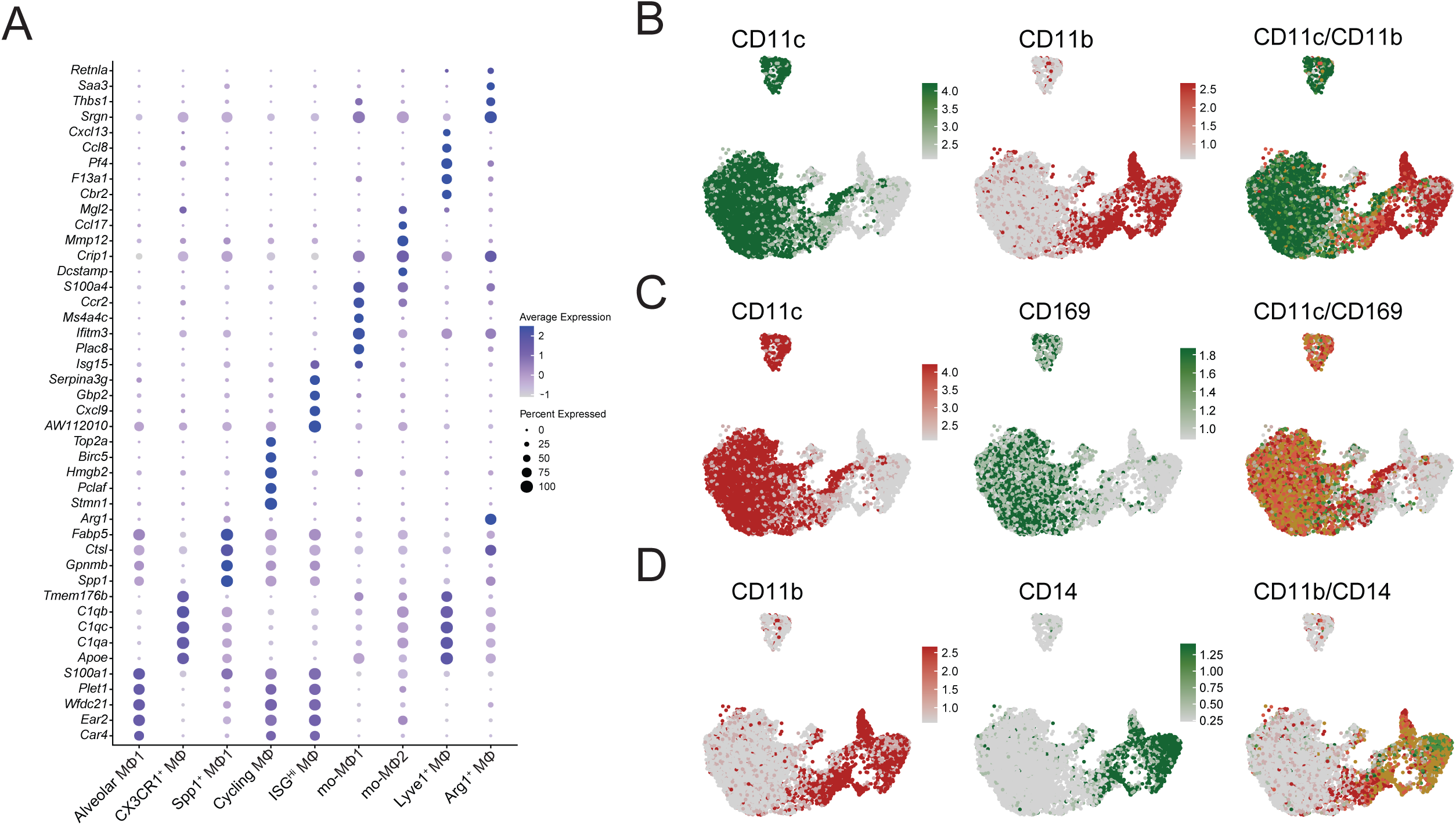
Macrophage subclusters in *Lkb1*-mutated tumors using ExCITE-seq. (A) Differentially expressed genes of macrophage subclusters from ExCITE-seq dataset. (B, C, D) UMAP visualization of antibody derived tag expression in macrophages for (B) CD11c/CD11b, (C) CD11c/CD169 and (D) CD11b/CD14 to identify alveolar and interstitial macrophages.

**Figure S9:**
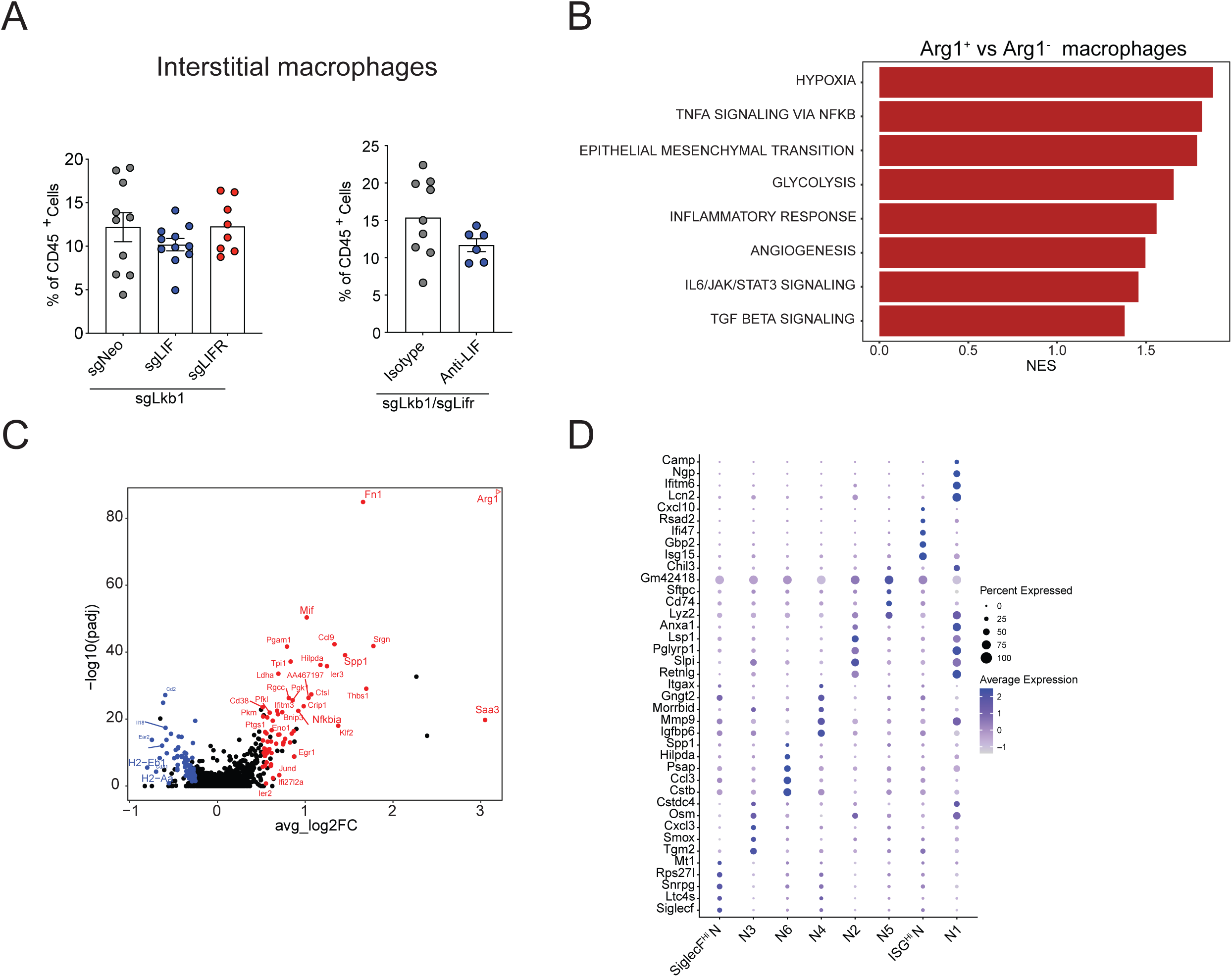
Ablation of either LIF or LIFR alters the transcriptional programming of myeloid cells. (A) Quantification of interstitial macrophages represented as a percentage of total tissue infiltrating immune cells (CD45^+^ CD45-Cir^-^) in *Lkb1*-mutant tumors with *Lif/Lifr* KO (left panel) or treated with anti-LIF neutralizing antibody (right panel). (B) Top upregulated pathways (FDR< 0.25) in macrophages comparing *Arg1^+^* vs Arg1^-^ macrophages in *Lkb1* mutant condition. (C) Volcano plot showing differential gene expression between *Arg1^+^* vs Arg1^-^ macrophages in *Lkb1* mutant condition using ExCITE-Seq dataset. Up-regulated genes are highlighted in red and down-regulated genes highlighted blue. Statistical analysis is outlined in Materials and Methods. (D) Differentially expressed genes of neutrophil subclusters from ExCITE-seq dataset. Individual mice are shown in the bar plots with mean and standard error. Statistical analysis was performed using Mann Whitney U or one-way ANOVA and Tukey’s test where appropriate. * p< 0.05 ** p<0.01 *** p<0.001 **** p< 0.0001

**Figure S10:**
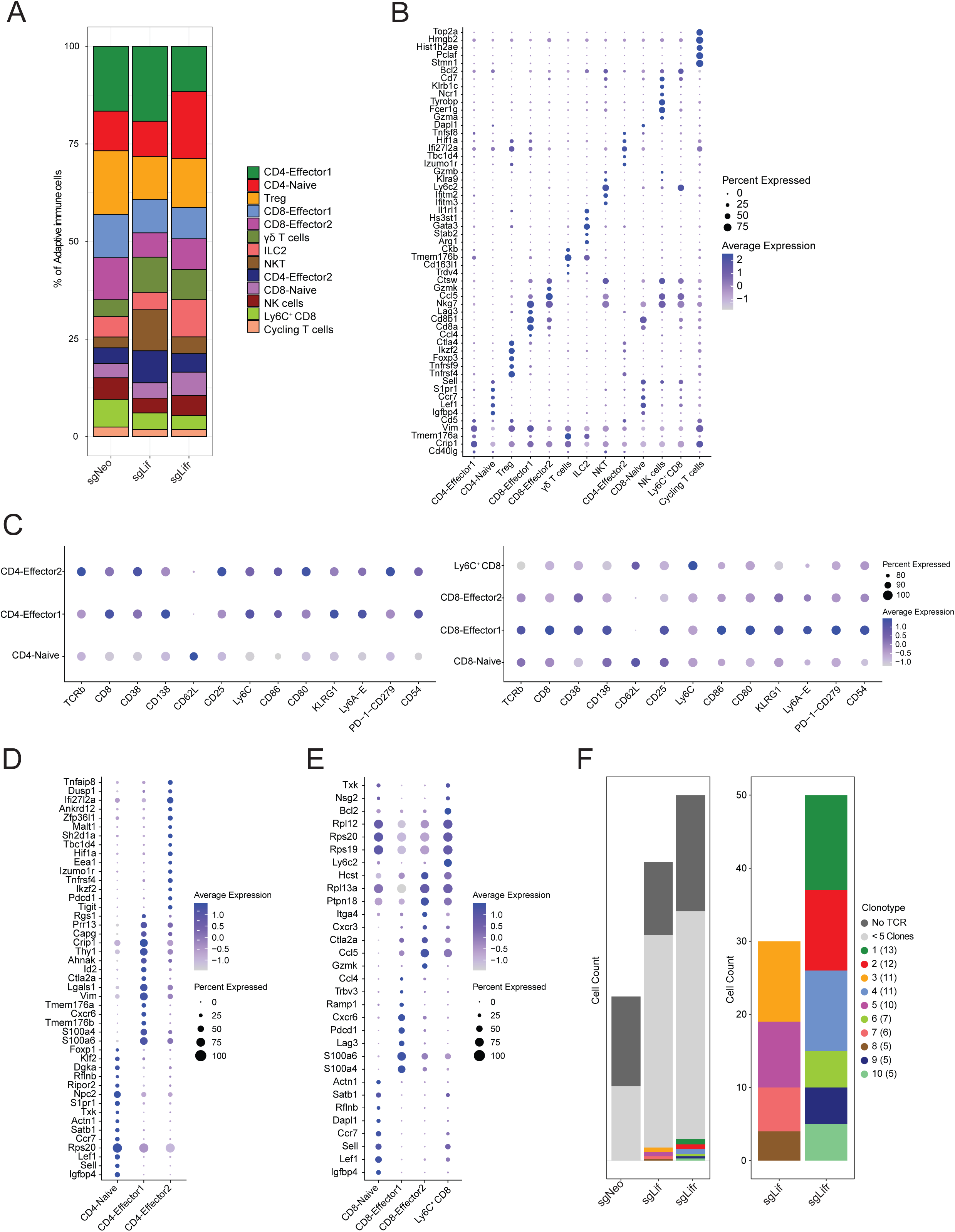
ExCITE-seq reveals TCR expansion in LIF/LIFR KO tumors. (A) Proportion of adaptive immune cell subclusters in each condition from ExCITE-seq dataset. (B) Top differentially expressed genes of adaptive immune cell subclusters. (C) Dot plot of antibody derived tag expression in CD4 and CD8 T cell subclusters for selected proteins. (D) Dot plot showing top differentially expressed genes of CD4 T cell subclusters. (E) Dot plot showing top differentially expressed genes of CD8 T cell subclusters. (F) The top ten most abundant clones with 5 cells or more for each condition are shown. Each bar is colored by individual clonotype with cell numbers for each clone included.

## Notes

### Competing Interest Statement

The authors have declared no competing interest.

